# Identification of functional non-coding variants associated with orofacial cleft

**DOI:** 10.1101/2024.06.01.596914

**Authors:** Priyanka Kumari, Ryan Z. Friedman, Lira Pi, Sarah W. Curtis, Kitt Paraiso, Axel Visel, Lindsey Rhea, Martine Dunnwald, Anjali P. Patni, Daniel Mar, Karol Bomsztyk, Julie Mathieu, Hannele Ruohola-Baker, Elizabeth J. Leslie, Michael A. White, Barak A. Cohen, Robert A. Cornell

## Abstract

Oral facial cleft (OFC) is a multifactorial disorder that can present as a cleft lip with or without cleft palate (CL/P) or a cleft palate only. Genome wide association studies (GWAS) of isolated OFC have identified common single nucleotide polymorphisms (SNPs) at the 1q32/*IRF6* locus and many other loci where, like *IRF6*, the presumed OFC-relevant gene is expressed in embryonic oral epithelium. To identify the functional subset of SNPs at eight such loci we conducted a massively parallel reporter assay in a cell line derived from fetal oral epithelium, revealing SNPs with allele-specific effects on enhancer activity. We filtered these against chromatin-mark evidence of enhancers in relevant cell types or tissues, and then tested a subset in traditional reporter assays, yielding six candidates for functional SNPs in five loci (1q32/*IRF6*, 3q28/*TP63*, 6p24.3/*TFAP2A*, 20q12/*MAFB*, and 9q22.33/*FOXE1*). We further tested two SNPs near *IRF6* and one near *FOXE1* by engineering the genome of induced pluripotent stem cells, differentiating the cells into embryonic oral epithelium, and measuring expression of *IRF6* or *FOXE1* and binding of transcription factors; the results strongly supported their candidacy. Conditional analyses of a meta-analysis of GWAS suggest that the two functional SNPs near *IRF6* account for the majority of risk for CL/P associated with variation at this locus. This study connects genetic variation associated with orofacial cleft to mechanisms of pathogenesis.

## Introduction

Oral facial cleft (OFC) is a multifactorial disorder that can present as a cleft lip with or without cleft palate (CL/P) or a cleft palate only (CP) and has a genetic predisposition. More than one hundred syndromes include OFC as a phenotype and overall about 30% of CL/P cases are syndromic^1^. Such syndromes are generally caused by mutations in single genes and follow a Mendelian inheritance pattern. The remaining cases are non-syndromic or isolated. The etiology of non-syndromic OFC is partially genetic as the concordance of non-syndromic OFC is 50% in identical twins but just 3-5% in other first-degree relatives^2–4^. Multiple independent genome wide association studies (GWAS), and meta-analyses thereof, have identified more than 40 loci where alleles of common single nucleotide polymorphisms (SNPs) are over-represented in cases versus in controls with the same ancestry (reviewed in^5^). Importantly, however, most of the heritable risk for isolated OFC has not been assigned to any gene or locus. Moreover, GWAS results alone do not illuminate the mechanisms of pathogenesis attributable to genetic variation at each locus. Understanding these mechanisms may guide the design of preventative therapies, and point to additional genes in which mutations will contribute risk for non-syndromic OFC.

Identifying functional (causal) SNPs among those in linkage disequilibrium with them is challenging. Most SNPs lie in non-coding DNA and the functional subset of them presumably disrupt *cis*-regulatory sequences (i.e., enhancers and promoters). However, the sequence constraints of *cis*-regulatory sequences remain poorly understood. We and others have used machine learning to mine sets of tissue-specific enhancers for sequence patterns^6–8^, and there are *in silico* tools for inferring variants that affect enhancer function^9, 10^, but currently there is no tool that can robustly identify non-coding variants that alter enhancer activity. An alternative approach is the massively parallel reporter assay (MPRA). MPRAs have been widely used to detect elements with enhancer activity, and in some cases to detect the effect of variants on enhancer activity^11–20^. A challenge in deploying MPRA to identify functional SNPs is that the results are highly dependent on the cellular context^15^ and so it is important to find a cell line that models the embryonic cell type in which the SNPs affect disease risk.

The 1q32/*IRF6* locus is associated with OFC in multiple ethnic groups^21–32^. *IRF6,* encoding the transcription factor Interferon Regulatory Factor 6, is strong candidate for the risk-relevant gene (i.e., the effector gene) in this locus because mutations in *IRF6* are found in about 70% of patients with Van der Woude syndrome, the most common syndromic form of OFC (VWS1, OMIM # 119300). Finding the functional SNPs at this locus can help uncover the gene regulatory networks governing morphogenesis of the face and identify new candidate genes for OFC. However, a meta-analysis of several GWAS identified more than 600 SNPs with *P* values indicating at least a suggestive association with OFC^23^. One of these SNPs, rs642961, resides in an enhancer of *IRF6* and altered the binding of the transcription factor AP2-α in an electrophoretic mobility shift assay, suggesting it was functional^33^. However, this conclusion was uncertain because in two studies this SNP did not have allele-specific effects in a standard reporter assay^33^ or in an MPRA^34^. The large number of OFC-associated SNPs at this locus makes it difficult to determine which are functional.

Here we deployed an MPRA in a fetal oral epithelium cell line to nominate candidate functional SNPs among those associated with OFC and within loci where the presumed effector gene is expressed in oral epithelium. We validated a subset of the MPRA results using traditional luciferase reporter assays in the cell line and in primary keratinocytes. For two promising SNPs near *IRF6* and one near *FOXE1*, we engineered the genotype of the SNPs in induced pluripotent stem cells, differentiated the cells into embryonic oral epithelium, and then assessed allele-specific effects on gene expression and transcription factor binding. These studies support six SNPs as being functional, with varying level of support, and show that two may account for most of the heritable risk for CL/P phenotype attributed to the *IRF6* locus in the cohort analyzed here.

## Results

### A massively parallel reporter assay reveals candidate functional SNPs at eight OFC-associated loci

Identifying the effector gene within GWAS-of-OFC loci is a challenge. For analysis by MPRA, here we picked 887 SNPs from eight such loci (1q32/*IRF6*, 2p21/*THADA,* 3q28/*TP63*, 6p24.3/*TFAP2A*, 9q22.2/*GADD45G*, 12q13.13/*KRT18*, 20q12/*MAFB*, and 9q22.33/*FOXE1*), in which the currently presumed effector gene is expressed in oral epithelium, although not necessarily only there, and which regulates differentiation of an epithelial tissue^35–60^ (Table 1, Supplementary Table 1, 2). At seven of the loci we picked SNPs based on their suggestive significance in a newly-performed meta-analysis of GWA studies (Table 1, Supplementary Table 2)^23^; we additionally included SNPs at the *TFAP2A* locus identified in an independent GWAS of CL/P in Han Chinese^26^. Because of constraints on the library size, for one locus (9q22.33/*FOXE1*) we instead picked SNPs in strong linkage disequilibrium with the lead SNP (rs12347191) in a European population and annotated by Haploreg as being within regulatory elements^23^ (Table 1, Supplementary Table 2). As outlined in Fig. 1a, we synthesized a library of reporter plasmids containing 161 base-pair (bp) genomic elements each centered on an OFC-associated SNP; OFC risk and non-risk alleles of each SNP were represented in four replicate constructs with distinct barcodes (Supplementary Table 3, 4). We performed the MPRA in GMSM-K cells (Supplementary Table 5), a cell line derived from human fetal oral mucosa^61^, and the results were strongly correlated across four replicate experiments (Supplementary Fig. 1a). The enhancer activities of the tested chromatin elements were modestly correlated with levels of histone H3 lysine 27 acetylation (H3K27Ac), a mark of active enhancers, in primary normal human epidermal keratinocytes (NHEKs) (Pearson correlation coefficient r = 0.02) (Supplementary Fig. 1b, Supplementary Table 6). Unexpectedly, the enhancer activity of four tested elements from within an element known as “multi species conserved sequence 9.7 kb from *IRF6*” (*IRF6* MCS9.7),^33^ which has enhancer activity in murine embryonic oral epithelium^62^ and in zebrafish periderm^6^, was only marginally higher than that of the average of the 84 scrambled elements (Supplementary Fig. 1c, Supplementary Tables 6, 7), probably because 161 bp captures only a portion of endogenous enhancers. Nonetheless, of the 887 SNPs tested, 65 had allele-specific effects on enhancer activity (Fig. 1b, Supplementary Table 6), nominating these as candidates for functional SNPs. There was at least one such candidate at each of the loci included in the study; at the 1q32/*IRF6* locus there were 46 such candidates (Fig. 1b and Table 1, Supplementary Table 6).

**Fig. 1:**
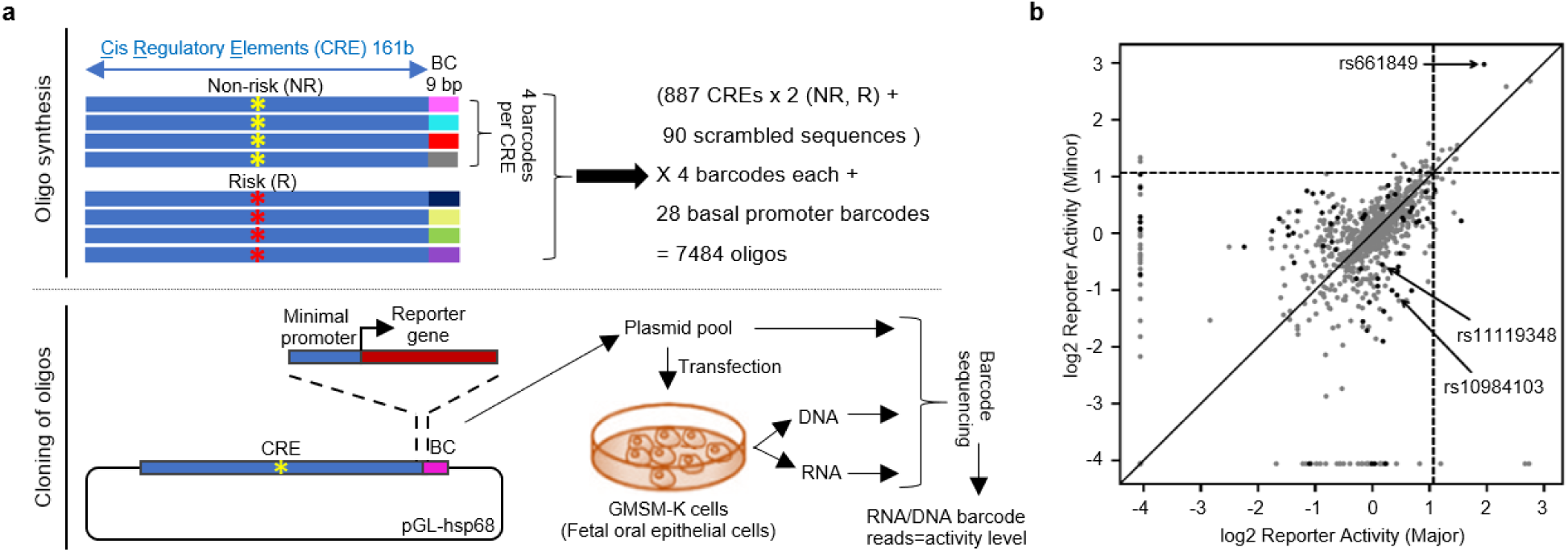
MPRA identified SNPs with allele-specific significant effects on reporter activity. **a)** Schematic of MPRA library construction and execution. BC, barcode. Yellow and red asterisks: non-risk and risk allele, respectively. **b)** Scatter dot plot showing (black dots) 65 SNPs with significant allele-specific effects on reporter activity in the MPRA and (gray dots) 822 SNPs without them. Arrows indicate the functional SNPs identified in this study (two near *IRF6 -* rs11119348 & rs661849 and one near *FOXE1*-rs10984103). Dashed lines indicate the 95^th^ percentile of the reporter activity of scrambled elements; on both axes, zero, i.e., log2(1) represents the reporter activity of the empty vector.

**Table 1:**
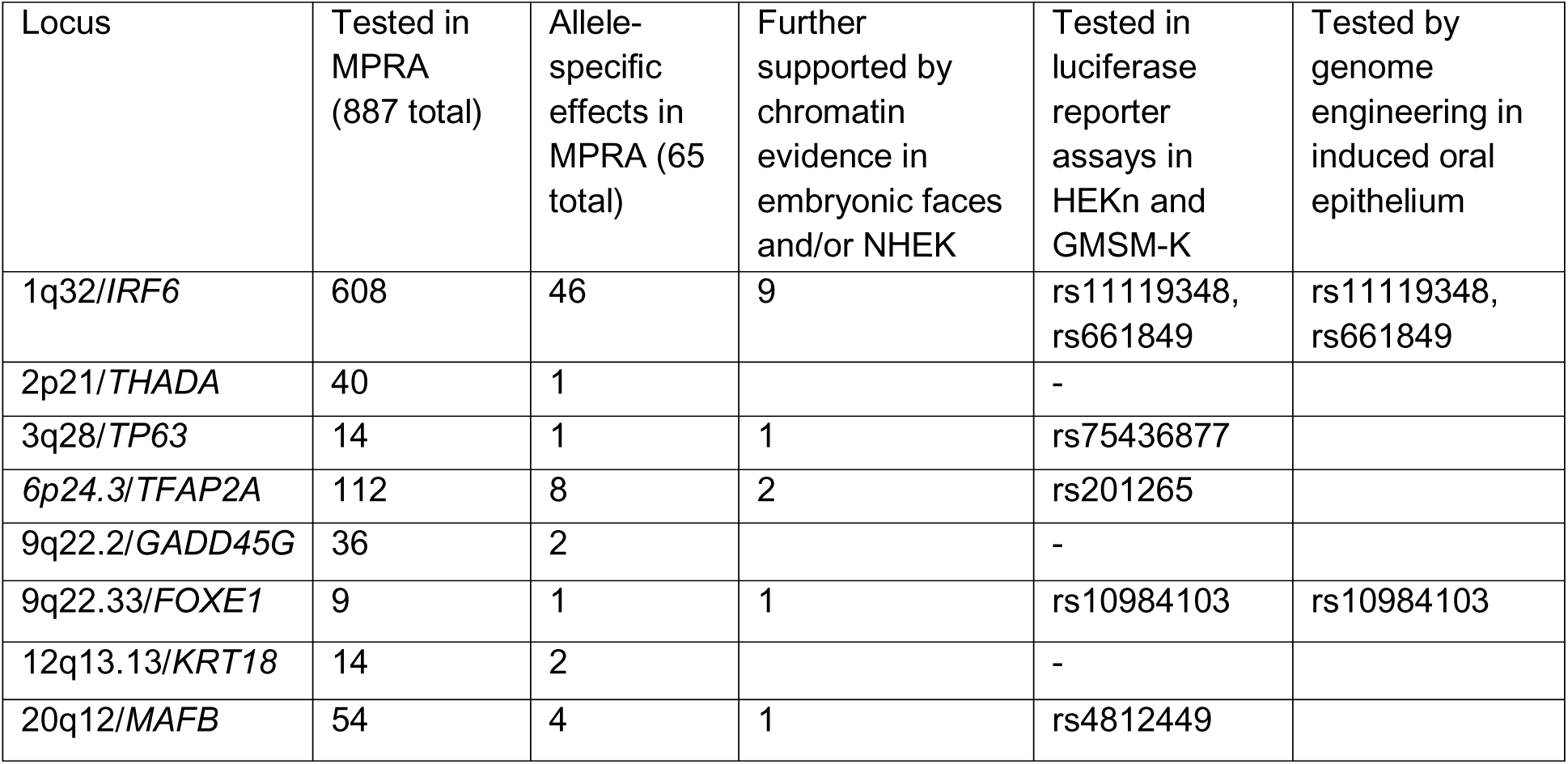
The number of SNPs tested, and the number with allele-specific effects, in the MPRA at each locus.

### Filtering candidate SNPs for those present in annotated enhancers

We next sought to filter MPRA-nominated SNPs for those lying within enhancers active in embryonic oral epithelium. In one approach to identify such enhancers, we used NHEKs as a model for embryonic oral epithelium and the activity-by-contact (ABC) method to identify enhancers for the presumed effector genes^63^. This method incorporates cell-type specific RNA-seq, DNAse hypersensitivity, and H3K27Ac ChIP-seq datasets, all of which are available for NHEKs, and averaged Hi-C chromatin-contact data from 10 ENCODE cell types^63, 64^. Within two apparent enhancers for *IRF6* and one for *FOXE1* there were MPRA-nominated SNPs (i.e., rs11119348 and rs661849 near *IRF6*, rs10984103 near *FOXE1*) (Fig. 2a-c). The first candidate SNP near *IRF6*, rs11119348, lies within *IRF6* MCS9.7. This element is the site of a rare single-base-pair duplication that appears to cause Van der Woude syndrome^65^. It also contains three other common SNPs (i.e., in addition to rs11119348) that are associated with non-syndromic OFC, including rs642961, the focus of an earlier study^33^. We tested all four of these common SNPs (separately) in the MPRA but only rs11119348 had allele-specific effects in it. We will refer to this as the *“IRF6 -*10 kb SNP” (Fig. 2b). The second SNP in this locus, rs661849, resides within an evolutionarily conserved sequence 21.7 kb upstream of the *IRF6* promoter and within the *UTP25* promoter. This SNP is an eQTL for *IRF6* in the brain (GTEx). There are no other OFC-associated SNPs within the same apparent regulatory element. We will refer to this SNP as the “*IRF6* -22 kb SNP” (Fig. 2b). Of note, rs642961, mentioned above, is in linkage disequilibrium with *IRF6* -22 kb SNP (R^2^ = 0.76) but not with *IRF6* -10 kb SNP (R^2^ = 0.03) (based on data from the 1000 Genomes project) (Supplementary Fig. 2 and Supplementary Table 8). The candidate SNP within a *FOXE1* enhancer, rs10984103, lies within a predicted enhancer 23.8 kb downstream of the *FOXE1* transcription start site (Fig. 2c). This SNP is an eQTL for *FOXE1* in multiple tissues including the brain (GTEx). We will refer to this SNP as the “*FOXE1* 24 kb SNP.”

**Fig. 2:**
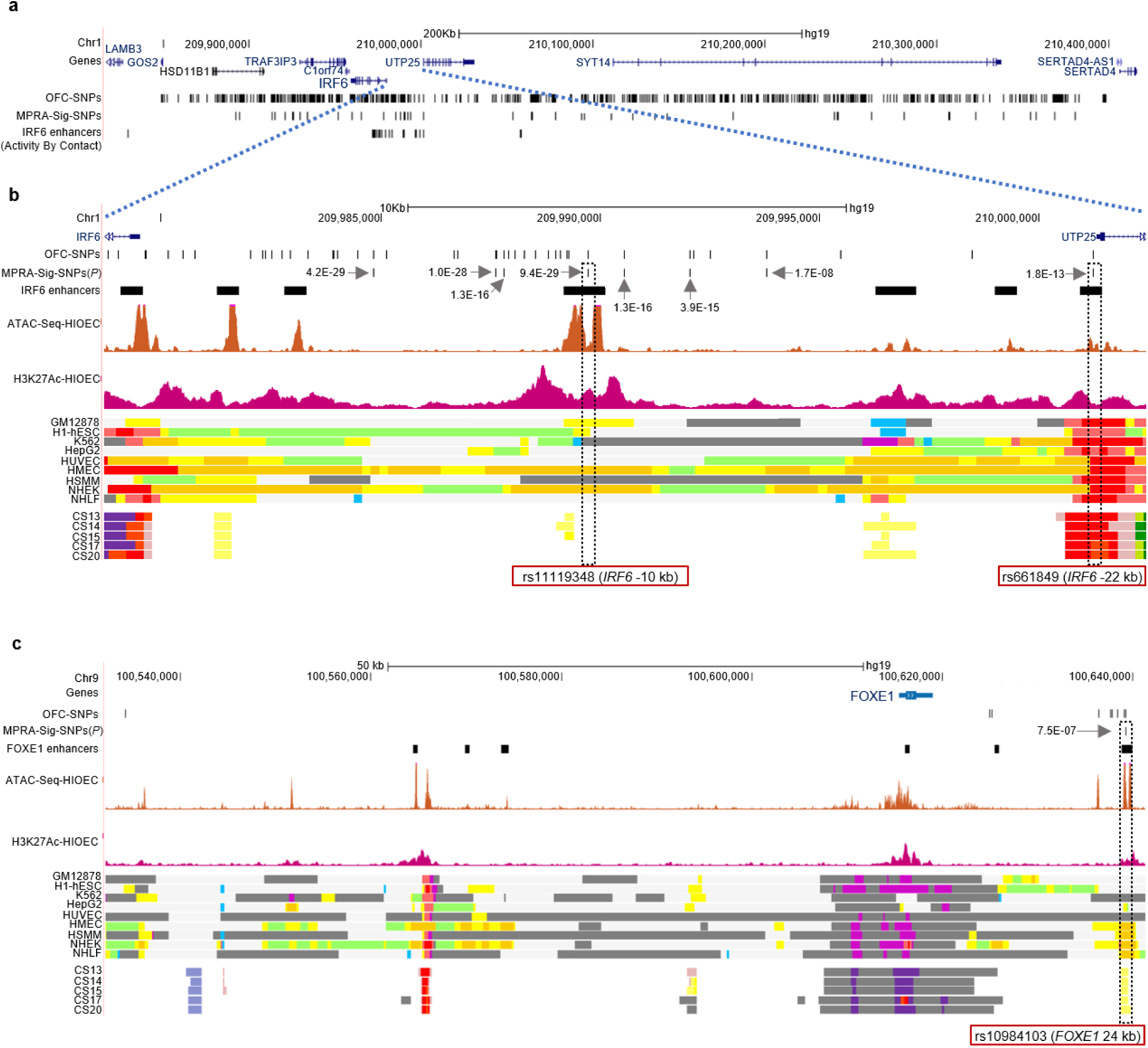
Two SNPs (rs11119348 and rs661849) at *IRF6* and one SNP (rs10984103) at *FOXE1* as regulatory variants. **a-c)** UCSC browser views of the human genome, GRCh37/hg19, focused on the OFC associated SNPs at **a, b**) *IRF6* and **c**) *FOXE1*. **OFC SNPs**, SNPs identified by GWAS and evaluated by MPRA at each locus. **MPRA-sig SNPS**, SNPs with allele-specific effects in the MPRA. ***IRF6* enhancers, *FOXE1* enhancers**, enhancers for the indicated gene identified using the Activity-by-Contact (ABC) method and datasets from NHEK (see Methods). **Gray arrows**, GWAS-meta-analysis *P* values of the indicated SNPs. **GM12878**, B-cell derived cell line; **H1-ESC**, embryonic stem cells; **K562**, myelogenous leukemia cell line; **HepG2**, liver cancer cell line; **HUVEC**, human umbilical vein endothelial cells; **HMEC**, human mammary epithelial cells; **HSMM**, human skeletal muscle myoblasts; **NHEK**, normal human epidermal keratinocytes; **NHLF**, normal human lung fibroblasts. **CS 13-20**, Carnegie stages for human embryonic face explants^68^. **Colored bars,** inferred chromatin state from combinatorial analysis of multiple chromatin mark datasets^106^. **Orange** and **yellow**, active and weak enhancer element, respectively; **bright red**, active promoter; **light red**, weak promoter; **purple**, inactive/poised promoter; **blue**, insulator; **light green**, weak transcribed; **grey**, Polycomb repressed; **light grey**, heterochromatin/repetitive; **green** and **dark purple** in facial explants dataset, transcription and bivalent promoter, respectively.

In a second, broader approach to identify potential embryonic oral epithelium enhancers that is agnostic to the identity of the effector gene we used chromatin-mark evidence of enhancers active in NHEK (ENCODE), in human immortalized oral epithelial cells (HIOEC), which are potentially a better model of embryonic oral epithelium than NHEK^66, 67^, or in human embryonic faces^68^. Within elements marked as enhancers in one or more of these contexts there were fourteen MPRA-nominated SNPs, including the three discussed above; the additional SNPs were detected in the 1q32/*IRF6*, 6p24.3/*TFAP2A,* 3q28/*TP63* and 20q12/*MAFB* loci (Supplementary Table 9, Supplementary Figs. 3-9). In summary, intersecting MPRA-nominated SNPs with predicted enhancers strengthened the candidacy of fourteen SNPs as being functional (Table 1, Supplementary Table 9).

### MPRA results tested with luciferase reporter assays

We next tested a subset of the MPRA results in traditional luciferase reporter assays in GMSM-K cells, using reporter elements arbitrarily chosen to be 701 bp long and centered on the SNP (Supplementary Tables 10, 11a). For these tests we picked six SNPs with allele-specific effects, and seven SNPs without them, in the MPRA. The former were all among the fourteen candidates discussed above and included *IRF6* -10 kb, *IRF6* -22 kb, *FOXE1* 24kb, rs4812449 (near *MAFB*), rs201265 (near *TFAP2A*), and rs75436877 (near *TP63*); the latter included rs642961 within *IRF6 MCS9.7* (Supplementary Tables 10, 11a). All six SNPs with allele-specific effects in the MPRA also had them in the luciferase reporter assays (Supplementary Fig. 10a and Supplementary Table 11a), although in two cases, *FOXE1* 24 kb and rs201265 (near *TFAP2A*), the direction of the effect was reversed (Supplementary Fig. 10a and Supplementary Table 11b). Discordance between the results from traditional reporter assays and MPRA in the direction-of-effects of SNP alleles has been reported in other studies^12, 69^ and presumably reflects that the short elements used in MPRAs are not entire enhancers (addressed below).

Among the seven SNPs that did not have allele-specific effects in the MPRA, five also did not have them in the luciferase reporter assays. These five included rs642961 within *IRF6 MCS9.7,* in agreement with a previous report^33^ (Supplementary Fig. 10a and Supplementary Table 11a). In summary, for eleven of thirteen SNPs tested, luciferase reporter assays agreed with MPRA regarding whether enhancer activity was affected by a SNP allele; this rate of concordance between luciferase and MPRA results is significant (*P* = 0.0163, one-sided Fisher’s exact test) and matches or exceeds that of other studies using MPRAs to detect functional non-coding SNPs^12, 13^.

We found that the length of the element tested, and whether it matched the extent of open chromatin of an endogenous enhancer, affected a SNP’s direction of effect in reporter assays. First, for the two SNPs with opposite directions of effect in MPRA and luciferase reporter assays (*FOXE1* 24 kb and rs201265), we repeated the latter using shorter elements that matched the lengths used in the MPRA (i.e., 161 bp). In these luciferase reporter assays, the results concorded with those from the MPRA (Supplementary Fig. 10b and Supplementary Table 11b). Second, for three promising SNPs (*IRF6* -10 kb, *IRF6* -22 kb and *FOXE1* 24 kb) with significant allele-specific effects in the MPRA and in the luciferase reporter assays, we repeated the latter using slightly longer elements (936 bp to 1.1 kb) that aligned with open chromatin (i.e., ATAC-seq peaks) in HIOEC and in NHEK (Supplementary Fig. 10c-e, Supplementary Table 10).

Remarkably, for all three SNPs the directions of effect were reversed relative to those using 701 bp elements: the risk-alleles of *IRF6* -10 kb and *IRF6* -22 kb SNPs reduced the activity, and the risk allele of *FOXE1* 24 kb elevated the activity, of the enhancers in which they reside (Supplementary Fig. 10c-e). The SNPs’ effect in reporter assays using the longest elements seemed most likely to reflect these effects in the endogenous enhancers; we test this prediction for three SNPs later in the study. To summarize, in reporter assays the length of the element tested affected a SNP’s direction of effect but not whether there was an effect.

At this point in the study we conducted RNA-seq on our stock of GMSM-K and discovered that, relative to primary adult keratinocytes (NHEK), GMSM-K express higher levels of *PITX1* and *FOXE1* transcripts, consistent with their origin in the oral cavity; however, unexpectedly, they expressed lower levels of epithelial markers *IRF6*, *CDH1, KRT8*, *KRT18* and *TP63* (Supplementary Fig. 11a, b and Supplementary Table 12). Therefore, we repeated the luciferase reporter assays in primary neonatal keratinocytes (HEKn), using the longer elements for the three SNPs where we had created them. Again, all six SNPs with allele-specific effects in the MPRA, and five of the seven SNPS without them, also had these qualities in luciferase reporter assays performed in HEKn cells (Fig. 3a-c, Supplementary Fig. 12). Thus, while our aliquot of GMSM-K lacks the robust epithelial features it was reported to have initially^61^, the results from an MPRA conducted in GMSM-K largely concorded with those from luciferase reporter studies in HEKn cells, which are epithelial.

**Fig. 3:**
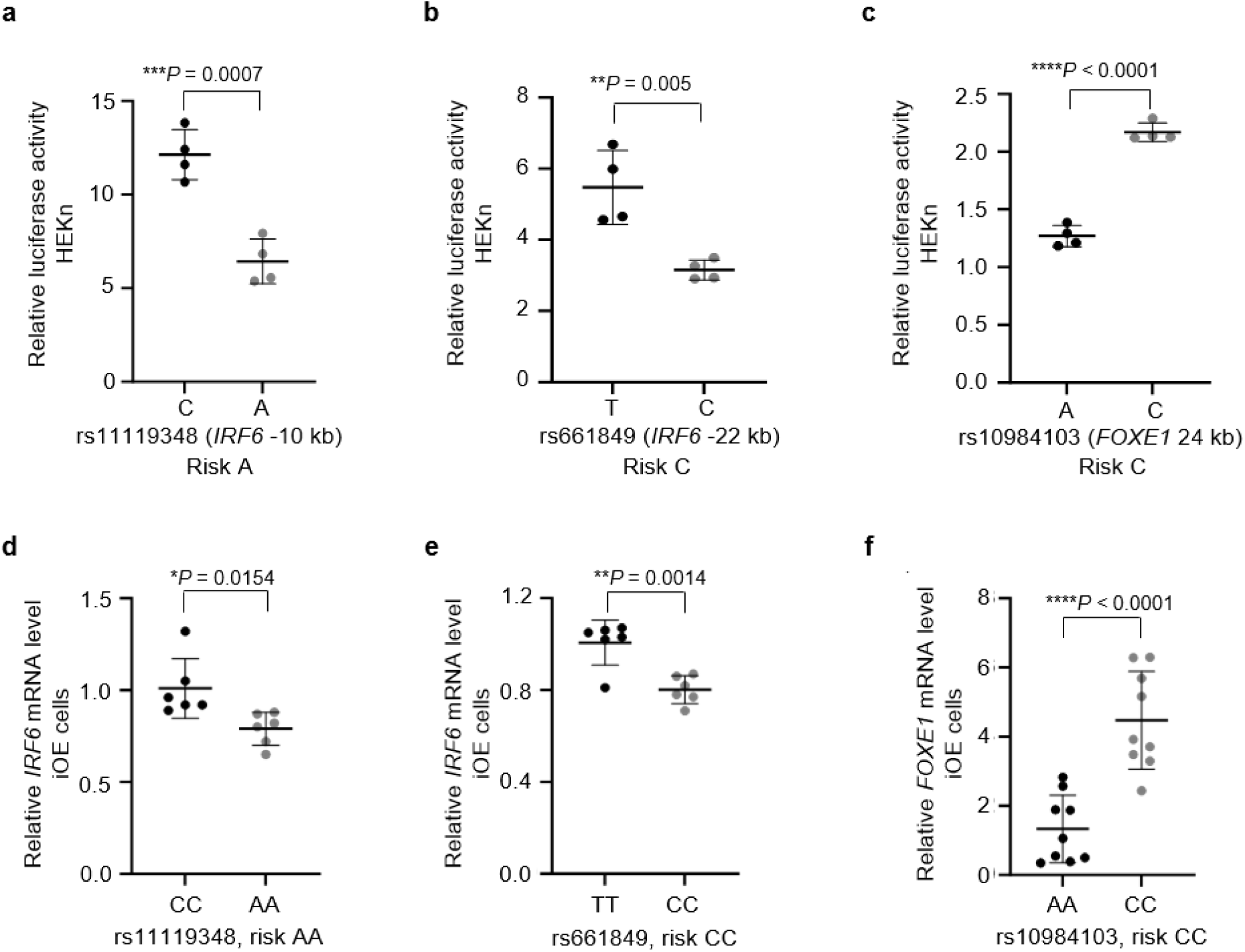
Risk alleles of two SNPs at *IRF6* and one SNP at *FOXE1* affect the enhancer activity and respective risk-associated gene expression. **a-c)** Scattered dot plot of relative luciferase activity (using the longer elements described in Results) for non-risk and risk alleles of rs11119348, rs661849 and rs10984103, respectively, in primary neonatal keratinocytes (HEKn). Value of 1 is that of the empty pGL3 promoter vector. Data are represented as mean ± standard deviation (SD) from four independent experiments. Statistical significance (*P* value, two-tailed) is determined by Student’s *t*-test. **d-f)** Scattered dot plot of relative levels of (d, e) *IRF6* or (f) *FOXE1* mRNA in edited iOE cells homozygous for the the non-risk or risk alleles of each SNP, as indicated, assessed by qRT-PCR. Expression levels are normalized against those of *ACTB, GAPDH, HPRT, UBC* and *CDH1*. Data are represented as mean ± SD from **(d, e)** six replicates or **(f)** nine replicates of cells harboring each genotype, as indicated in the plot. Statistical significance (*P* value, two-tailed) is determined by Student’s *t*-test.

### Genome engineering reveals rs11119348 and rs661849 affect *IRF6* expression, and rs10984103 affects *FOXE1* expression

To test whether the *IRF6* -10 kb and *IRF6* -22 kb SNPs affect enhancer activity in their native genomic contexts we engineered the genotype of WTC-11 induced pluripotent stem cells (iPSCs), which are heterozygous for the risk and non-risk alleles of both SNPs. For each SNP we raised two clones each that were homozygous for the risk allele or for the non-risk allele (Supplementary Figs. 13, 14). Next, we subjected three replicates of each clone to a 10-day protocol that induces iPSCs to differentiate into embryonic oral epithelial cells (induced oral epithelial cells, iOE)^70^ (Supplementary Fig. 13). Quantitative RT-PCR (qRT-PCR) revealed that the average level of *IRF6* expression in cells homozygous for the risk allele of either SNP (and heterozygous at the other) was lower than that in clones homozygous for the non-risk allele of that SNP (Fig. 3d, e), consistent with the luciferase reporter assays using the longest elements. We also engineered the genotype of these SNPs in GMSM-K. Despite the much lower expression of *IRF6* in GMSM-K compared to in iOE cells, cells homozygous for risk alleles had lower *IRF6* expression than cells homozygous for non-risk alleles (Supplementary Fig. 15a-f). These results indicate that the *IRF6* -10 kb SNP and the *IRF6* -22 kb SNP both directly affect *IRF6* expression.

Similarly, we engineered the genotype of the *FOXE1* 24 kb SNP. The parental iPSCs were heterozygous for the risk and non-risk alleles. We engineered three clones each that were homozygous for the risk allele or the non-risk allele and differentiated them in triplicate into iOE cells (Supplementary Fig. S16a, b). qRT-PCR revealed that the average level of *FOXE1* in cells homozygous for the risk allele was higher than that of the homozygous for the non-risk allele (Fig. 3f), consistent with the luciferase reporter assays using the longest elements, indicating that the *FOXE1* 24 kb SNP directly affects *FOXE1* expression.

### The *IRF6* -10 kb SNP risk allele promotes binding of the OFC-associated transcriptional repressor FOXE1

Using an online tool^71^ we predicted that the risk allele of the *IRF6* -10 kb SNP diminishes the affinity for two transcription factors (i.e., AR and SOX10) and elevated it for POU5F1B plus sixteen members of the FOX family, including FOXE1 (JASPAR score for FOXE1 site MA1487.1: risk, 14.1; non-risk, 8.6) (Fig. 4a, Supplementary Table 13a, b). Members of the FOX family can function as transcriptional activators or repressors^72, 73^. In a published report, chromatin immuno-precipitation with sequencing (ChIP-seq) revealed binding of FOXA1 and FOXA2 in bronchial epithelial cells at the *IRF6* -10 kb enhancer, and knocking down expression of either transcription factor led to upregulation of *IRF6*^74^. However, we found no published reports connecting FOXA1 or FOXA2 to craniofacial defects. By contrast, mutations in *FOXE1* cause Bamforth-Lazarus syndrome^75^ and SNPs near it are associated with risk for non-syndromic OFC^23, 76, 77^. Using an antibody against FOXE1 we conducted ChIP in iOE cells generated from the parental iPSCs. qPCR revealed more chromatin precipitated at the *IRF6* -10 kb SNP locus by anti-FOXE1, and by anti-H3K27Ac, than at an intergenic region, consistent with the SNP being in an enhancer bound by FOXE1 (Fig. 4b, c). While the iOE cells were heterozygous at this SNP (like the parental iPSCs) (Fig. 4d), PCR and sequencing revealed that chromatin precipitated by the anti-FOXE1 antibody was enriched for the risk allele (Fig. 4d, two additional replicates in Supplementary Fig. 17a). By contrast, the anti-H3K27Ac antibody pulled down more of the non-risk allele (Fig. 4d, Supplementary Fig. 17a), consistent with higher activity of the enhancer and higher expression of *IRF6* in cells with the non-risk genotype. Similarly, anti-FOXE1 ChIP in GMSM-K homozygous for the risk or non-risk allele of *IRF6* -10 kb SNP revealed stronger binding of FOXE1 to the risk allele (Supplementary Fig. 18a). These results indicate that the *IRF6* -10 kb SNP risk allele favors the binding of FOXE1.

**Fig. 4:**
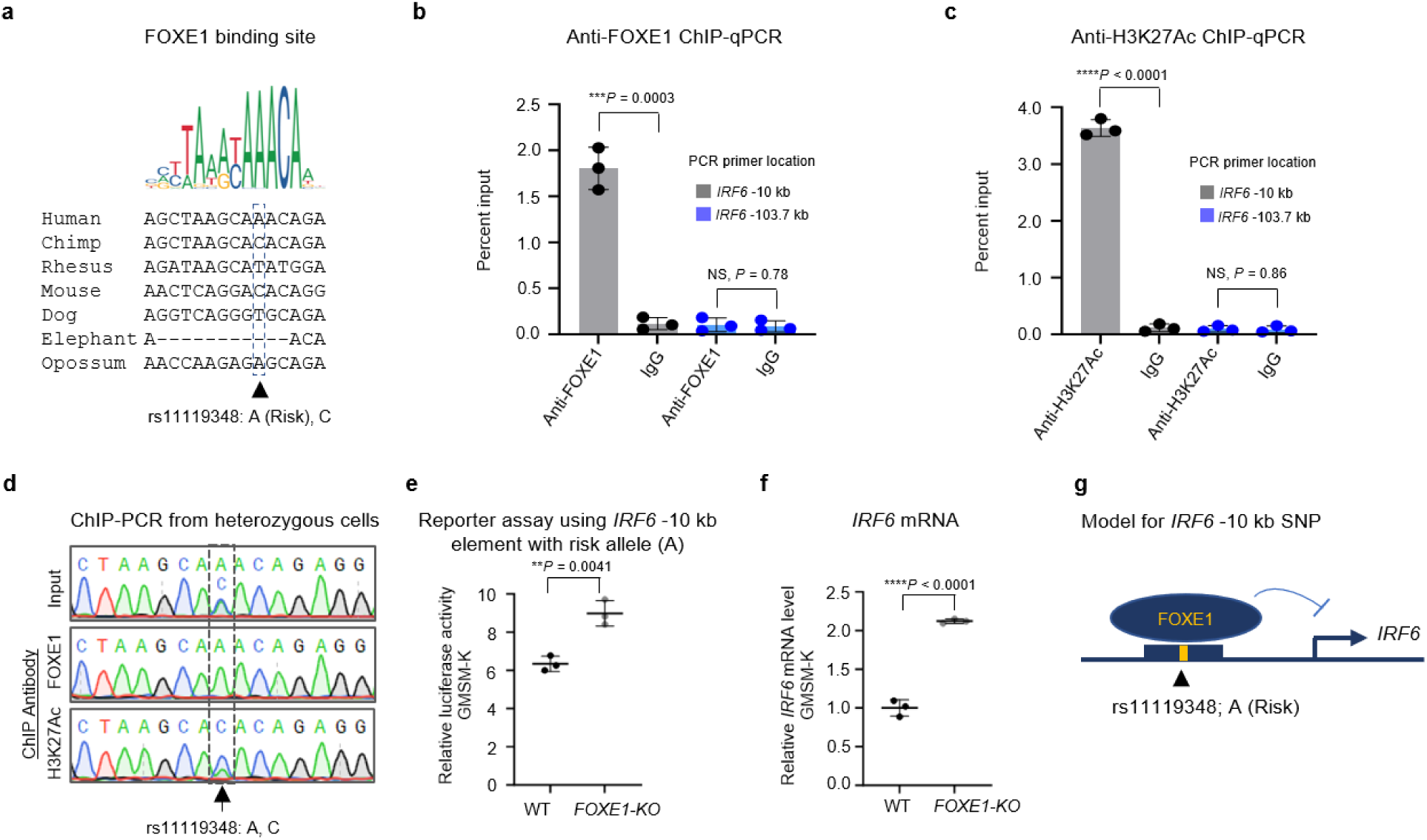
Risk allele of *IRF6* -10 kb SNP (rs11119348) promotes binding of transcriptional repressor FOXE1. **a)** Consensus FOXE1 binding motif from the JASPAR database of transcription factor DNA-binding preferences (Matrix ID: MA1847.1) and alignment of the variant site in several mammals. The risk allele (A) is the reference allele and has a higher frequency than the non-risk allele (C) in most populations. **b, c)** Percent input identified by ChIP-qPCR for anti-FOXE1 and anti-H3K27Ac, respectively, in iOE cells heterozygous for rs11119348 using primers specific to the *IRF6* -10 kb enhancer site or, as a negative control, to a region 103.7 kb upstream of the *IRF6* transcription start site that lacked ATAC-Seq and H3K27Ac ChIP-Seq signals in HIOEC or NHEK. Error bars refer to three ChIP replicates and expressed as mean ± SD. Statistical significance (*P* value, two-tailed) is determined by Student’s *t*-test. NS, non-significant. **d)** Sequencing of anti-FOXE1 and anti-H3K27Ac ChIP-PCR products from cells heterozygous for rs11119348 using the indicated antibody. **e)** Scattered dot plot of relative luciferase activity of the *IRF6* -10kb reporter construct (longest version) harboring the risk allele of rs11119348, in wildtype (WT) or *FOXE1*-KO GMSM-K cells. Data are represented as mean ± SD from three independent experiments. Statistical significance (*P* value, two-tailed) is determined by Student’s *t*-test. **f)** Scattered dot plot of relative levels of *IRF6* mRNA in WT and *FOXE1*-KO GMSM-K cells assessed by qRT-PCR. Expression levels of *IRF6* are normalized against *ACTB*. Data are represented as mean ± SD from three replicates. Statistical significance (*P* value, two-tailed) is determined by Student’s *t*-test. **g)** Model showing binding of FOXE1 to the *IRF6* -10 kb enhancer, which if favored by the risk allele, reducing *IRF6* expression.

To test the importance of FOXE1 in regulating the *IRF6* -10 kb enhancer we repeated the luciferase reporter assays using the *IRF6* -10 kb enhancer (longest version) harboring the risk allele of the *IRF6* -10 kb SNP, and compared the results between parental GMSM-K and those in which we had deleted the *FOXE1* using CRISPR/Cas9 reagents (Supplementary Figs. 18b, 19). We chose GMSM-K over HEKn because the expression of *FOXE1* is higher in GMSM-K than in HEKn (Supplementary Fig. 11c). Interestingly, in cells lacking *FOXE1* the activity of the *IRF6 -*10 kb reporter was higher than in the control cells (Fig. 4e). We saw analogous results in cells transfected with siRNA targeting *FOXE1* or a non-targeting siRNA (Supplementary Fig. 18c, d). Moreover, levels of *IRF6* expression were higher in *FOXE1*-knockout or *FOXE1* knockdown GMSM-K than in the respective controls (Fig. 4f, Supplementary Fig. 18e). While the binding of other transcription factors may be affected by the allele of the *IRF6* -10 kb SNP, these results suggest its effect on binding of FOXE1 is sufficient to alter the activity of this enhancer (Fig. 4g).

### The *IRF6* -22 kb SNP risk allele promotes binding of the transcriptional repressor ETS2

Similarly, we discovered that the risk allele of the *IRF6* -22 kb SNP diminished the predicted affinity for two transcription factors (GATA2 and HSF4) and elevated it for NFKB2, ZNF454, and ten members of the ETS family, including ETS2 (JASPAR score for ETS2 site MA1484.1, risk: 4.9; non-risk, -4.1) (Fig 5a, Supplementary Table 14a, b). ETS2 can function as a transcriptional repressor^78, 79^ and suppresses differentiation of epithelia^80^. Anti-ETS2 and Anti-H3K27Ac antibodies yielded higher ChIP-qPCR signals in iOE cells at the *IRF6* -22 kb SNP than at an intergenic region (Fig. 5b, c). In iOE cells that were heterozygous for risk and non-risk alleles of this SNP PCR and sequencing of chromatin precipitated by anti-ETS2 revealed an enrichment of ETS2 binding to the risk allele (Fig. 5d, two additional replicates in Supplementary Fig. 17b), while anti-H3K27Ac pulled down more of the non-risk allele (Fig. 5d, Supplementary Fig. 17b), consistent with higher activity of the enhancer and higher expression of *IRF6* in cells with the non-risk genotype. Similarly, anti-ETS2 ChIP in GMSM-K homozygous for the risk or non-risk allele of *IRF6* -22 kb SNP revealed stronger binding of ETS2 to the risk allele (Supplementary Fig. 18f).

**Fig. 5:**
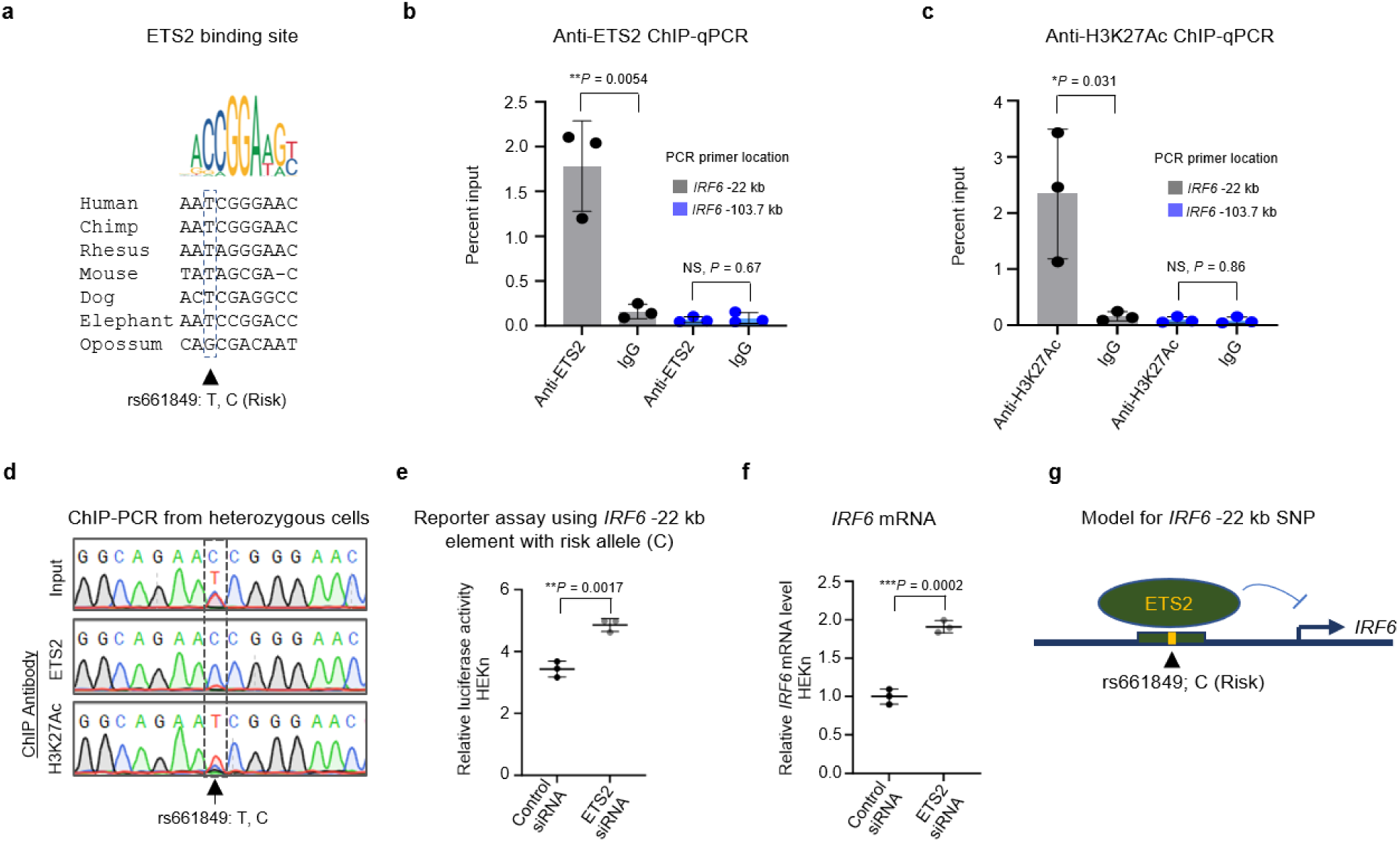
Risk allele of *IRF6* -22 kb SNP (rs661849) promotes binding of transcriptional repressor ETS2. **a)** Consensus ETS2 binding motif from the JASPAR database of transcription factor DNA-binding preferences (Matrix ID: MA1484.1) and alignment of the variant site in several mammals. The risk allele (C) is the reference allele but has a lower frequency than the non-risk allele (T) in most populations. The risk allele improves the match to the ETS2 binding site which remains partial. **b, c)** Percent input identified by ChIP-qPCR for anti-ETS2 and anti-H3K27Ac respectively in iOE cells heterozygous for rs661849 using primers specific to the *IRF6* -22 kb enhancer site or, as a negative control, to a region 103.7 kb upstream *IRF6* transcription start site lacking ATAC-Seq and H3K27Ac ChIP-Seq signals in HIOEC or NHEK. Error bars refer to three ChIP replicates and expressed as mean ± SD. Statistical significance (*P* value, two-tailed) is determined by Student’s *t*-test. NS, non-significant. **d)** Sequencing of anti-ETS2 and anti-H3K27Ac ChIP-PCR product of cells heterozygous for rs661849 using the indicated antibody. **e)** Scattered dot plot of relative luciferase activity of the *IRF6* -22kb reporter construct (longest version) harboring the risk allele of rs661849 in control versus *ETS2*-depleted primary neonatal keratinocytes (HEKn). Data are represented as mean ± SD from three independent experiments. Statistical significance (*P* value, two-tailed) is determined by Student’s *t*-test. **f)** Scattered dot plot of relative levels of *IRF6* mRNA in control versus *ETS2*-depleted HEKn assessed by qRT-PCR. Expression levels of *IRF6* are normalized against *ACTB*. Data are represented as mean ± SD from three replicates. Statistical significance (*P* value, two-tailed) is determined by Student’s *t*-test. **g)** Model showing binding of ETS2 to the *IRF6* -22 kb enhancer, which if favored by the risk allele, reducing *IRF6* expression.

We also found that reporter activity of the *IRF6* -22 kb enhancer harboring the risk allele of the *IRF6* -22 kb SNP was higher in ETS2-depleted HEKn relative to those transfected with a control siRNA (Fig. 5e, Supplementary Fig. 18g); we observed analogous results in GMSM-K (Supplementary Fig. 18h, i). *IRF6* expression was also higher in ETS2-depleted HEKn or GMSM-K than in control transfected cells (Fig. 5f, Supplementary Fig. 18j). Together these results support the model that the risk allele of the *IRF6* -22 kb SNP results in lower expression of *IRF6* by promoting the binding of ETS2, potentially among effects on the binding of other transcription factors (Fig. 5g, Supplementary Table 14a, b).

Similarly, we observed that the risk allele of the *FOXE1* 24 kb SNP diminished the predicted affinity for four transcription factors (GATA2, FOXH1, GCM2, and NRL) and elevated it for three transcription factors (TFAP2A, SP2, and KLF12) (Supplementary Table 15a, b). While we did not attempt chromatin immunoprecipitation (ChIP) with antibodies to transcription factors at this locus, ChIP with an anti-H3K27Ac antibody precipitated more of the risk allele of the *FOXE1* 24 kb SNP from heterozygous iOE cells (Supplementary Fig. 20a-c), consistent with higher activity of the enhancer and higher expression of *FOXE1* in cells with the risk genotype.

### Conditional analysis of haplotypes containing *IRF6* -10 kb and *IRF6* -22 kb SNP

Finally, to assess whether the *IRF6* -10 kb (rs11119348) and *IRF6* -22 kb (rs661849) SNPs can explain the heritable risk for CL/P associated with 1p22 we conducted conditional analyses by comparing cleft lip only (CL) and cleft lip and palate (CLP) cases separately to unrelated controls (we lacked sufficient cleft-palate-only cases to do a conditional analysis) (Fig. 6, Supplementary Table 16, 17). As expected, the *IRF6* locus was associated with CL (lead SNP rs12403599, *P* value = 3.29×10^−9^) (Fig. 6a) and CLP (lead SNP rs2076149, *P* value = 5.24×10^− 10^) (Fig. 6e). When compared to the unconditioned analysis, both *IRF6* -22 kb and rs11119348 explained some of the signal at the *IRF6* locus. However, *IRF6* -22 kb explained more signal in CL cases than in CLP cases. In other words, when we fixed on an allele of *IRF6* -22 kb, there was a bigger change in the *P* value for CL (lead SNP rs12403599, *P* value = 7.48×10^−6^) than for CLP (lead SNP rs2076149 *P* value = 3.34×10^−8^) (Fig. 6b, f). Conversely, *IRF6* -10 kb explained more signal in CLP cases (lead SNP rs55645018, *P* value = 0.0001) than in CL cases (lead SNP rs12403599, *P* value = 5.05×10^−6^) (Fig. 6c, g). Analyses conditioned on both SNPs explained most of the signal at *IRF6*, especially in CLP cases (lead SNP rs12403599 *P* value in CL = 0.0003; lead SNP rs55645018 *P* value in CLP = 0.0001) (Fig. 6d, h). The haplotypes containing the *IRF6* -10 kb and *IRF6* -22 kb SNPs together appear to explain most of the risk for CLP in the cohort analyzed here.

**Fig. 6:**
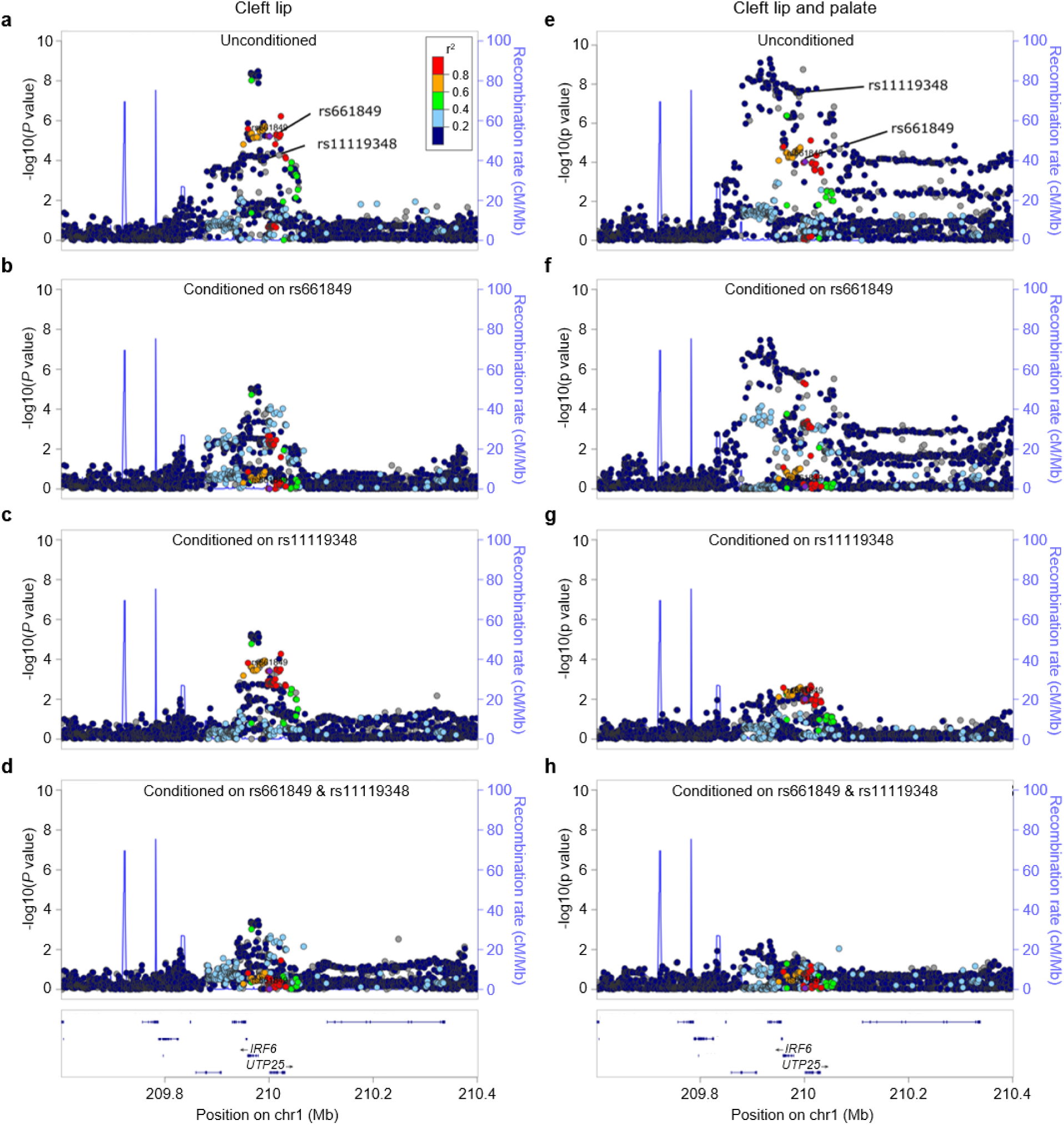
The haplotypes containing rs11119348 and rs661849 together explain much of the CL association, and most of the CLP association, in Europeans at the 1q32/*IRF6* locus. **a-h**) Locus Zoom plots of **a-d**) cleft lip only and **e-h**) cleft lip and palate; **a, e**) unconditioned; **b, f**) conditioned on rs661849 (*IRF6* -22 kb); **c, g**) conditioned on rs1119348 (*IRF6* -10 kb); **d, h**) conditioned on both SNPs simultaneously. Points are color-coded based on linkage disequilibrium (r^2^) in Europeans.

## Discussion

We sought to identify functional SNPs associated with OFC to understand how common variants predispose individuals to orofacial cleft (OFC). We tested 887 OFC-associated SNPs in a massively parallel reporter assay (MPRA) as has been done in other studies seeking to identify SNPs that affect enhancer activity^13, 81, 82^. Because enhancers are often cell-type specific, we limited the study to SNPs from loci where the presumed effector gene is expressed in oral epithelium (as opposed to oral mesenchyme or brain) and used a cell line model of this tissue for the MPRA. We then filtered SNPs with allele-specific effects in the MPRA for those that lie in enhancers active in a keratinocyte cell line (NHEK) and known to contact the promoter of the predicted risk gene in the locus. This yielded three top candidate SNPs, including two at *IRF6* and one at *FOXE1*. Filtering on MPRA-nominated SNPs on enhancers active in explants of human embryonic faces or in NHEK yielded additional candidate functional SNPs at *IRF6*, *TFAP2A, MAFB,* and *TP63.* We then used luciferase reporter assays to confirm the findings of the MPRA at several SNPs, including the three top candidate functional SNPs, and three additional candidates. Finally, we engineered the genome of iPSCs to be homozygous for the risk or non-risk allele of the three top-candidates SNPs (separately) and differentiated the cells into embryonic oral epithelium. We found that all three SNPs had allele-specific effects on expression levels of the respective effector gene, supporting the notion that these are bona fide functional SNPs. Risk alleles of both SNPs at *IRF6* decreased the expression level of *IRF6* whereas risk allele of the SNP at *FOXE1* increased the expression level of *FOXE1*. Both gain-of-function and loss-of-function mutations in *FOXE1* are cases with Bamforth–Lazarus Syndrome (which includes OFC)^41^, and over-expression of *Foxe1* can cause cleft palate in mice^46^. We identified transcription factors whose binding is affected by the SNP allele for both SNPs at *IRF6*, and showed that they regulate the expression of *IRF6*, and of enhancers harboring the SNPs, in the predicted direction. Finally, we returned to the GWAS data and found that the haplotypes containing the two SNPs explain most of the CLP association signal at this locus in the cohorts evaluated. In summary, we found strong evidence for three functional SNPs and suggestive evidence for three more.

This study illustrates how identifying functional SNPs at an OFC GWAS locus can illuminate transcriptional regulatory networks which in turn can identify additional OFC risk genes. For instance, analysis of transcription factor binding sites affected by the SNPs pointed to possible transcription factor families affected by each, and the *IRF6* -10 kb SNP affects a FOX binding site. We found that binding of FOXE1 is affected by the allele of this SNP; that of FOXA1 and FOXA2 may also be affected. We found that deletion of *FOXE1* elevates *IRF6* expression, which is consistent with FOXE1 binding to the *IRF6* -10 kb element and thereby dampening *IRF6* expression. *FOXE1* is implicated in risk for OFC by genome wide linkage analysis ^23, 77, 83^ and GWAS^26^, and mutations in it cause Bamforth-Lazarus syndrome that includes OFC ^84^.

These results support the possibility of a greater-than-additive effect on OFC risk in individuals harboring risk-associated haplotypes at both *IRF6* and *FOXE1*, but current GWAS data are underpowered to reveal such gene-by-gene interactions. Nonetheless, this example illustrates that identifying a functional SNP for OFC can indeed lead one to another OFC risk-relevant gene. In this light, it is interesting that the risk allele of the *IRF6* -22 kb SNP creates a low-affinity ETS family binding site, and ChIP-qPCR indicated that the risk allele promotes binding of ETS2. Low-affinity ETS binding sites are important for the regulation of SHH in the limb ^85^. In mice it is unclear if ETS2 governs craniofacial development because a strong loss-of-function mutation in *ETS2* causes embryonic lethality by E8.5^86^ *ETS2* is expressed in differentiated keratinocytes and glandular epithelium^87, 88^; it suppresses epithelial differentiation, and may contribute to carcinogenesis^80^. It would be interesting to test for genetic interactions between *Irf6* and *Foxe1*, or between *Irf6* and *Ets2*, in animal model organisms. In summary, *ETS2* and the genes encoding other transcription factors whose binding is predicted to be affected by the *IRF6* -22 kb SNP, *IRF6* -10kb, or *FOXE1* 24kb SNPs are worth exploring as OFC candidate genes.

The *IRF6* -10kb SNP is the fourth example of a potentially functional SNP associated with orofacial cleft within the same evolutionarily conserved enhancer of IRF6, MCS9.7^6, 33, 50, 62, 65, 89^. The first of these to be identified, rs642961, is associated with CL and the risk-associated allele disrupts binding of the transcription factor AP2-α (TFAP2A) in an electrophoretic mobility shift assay^33^. However, the present MPRA and a previous one^34^, and the current luciferase reporter assays and previous one^33^, all did not detect allele-specific effects of rs642961 on enhancer activity, suggesting it is not a functional SNP. It is possible that the *IRF6* -22kb SNP described here, or rs76145088 previously found to have allele-specific effects in an MPRA^34^, which are both in linkage disequilibrium with rs642961, are functional SNPs explaining the OFC signal leading investigators to rs642961. Subsequently, a rare, single nucleotide insertion in MCS9.7 was found to segregate with cleft palate and other characteristic symptoms in a Brazilian family with Van der Woude syndrome^65^. In stable transgenic mice, MCS9.7 (from the human genome) drove reporter expression in oral epithelium^62^; in stable transgenic zebrafish (*Tg*(*MCS9.7:gfp*), it drove reporter expression in periderm in the entire embryo including in the oral cavity^6^. We recently found that a SNP in the same enhancer, rs570516915, is associated with CP only in populations from Finland and Estonia and that the risk allele reduces *IRF6* expression level by disrupting the *IRF6* binding site^89^. The current study shows that rs11119348 in the same enhancer affects *IRF6* expression, possibly by altering the binding of FOXE1. Taken together, these studies emphasize the extraordinary importance of a single enhancer in genetic predisposition for orofacial cleft, and raise the question of how different mutations in this enhancer elevate risk for cleft lip with or without cleft palate, or cleft palate only.

There remains a need for high-throughput methods to screen functional SNPs in OFC and other common disorders. The MPRA we performed required synthesis of a library of oligos. This step was avoided in an MPRA that used reporter constructs from sheared chromatin from individual of distinct genotypes; this study managed to score more than 50 million SNPs^15^. However, the impact of this large scale MPRA was limited by only being performed in two cell types. The success of MPRAs at identifying functional SNPs relies on the cell line deployed being a reasonable model of the embryonic tissue where the SNP affects disease risk *in vivo*. While results from an MPRA helped us identify six OFC-associated SNPs that alter enhancer activity in luciferase reporter assays, we only pursued three of them with the in-depth experiments, confirming that they can alter *IRF6* or *FOXE1* expression in induced oral epithelium cells. It will be interesting to assess whether additional MPRA-nominated SNPs within active enhancers near *IRF6* are indeed functional, and if they affect *IRF6* expression in combination. However, the step of engineering the genome in iPSCs and creating induced oral epithelium or oral mesenchyme remains laborious and expensive. One promising approach to prioritize non-coding SNPs is chromatin-accessibility QTL analyses^90^ but these too require appropriate cell line models. Despite these challenges, identifying functional SNPs remains an essential step in translating the findings of GWAS into an understanding of how common genetic variants contribute to pathogenesis of OFC and other congenital anomalies.

## Materials and Methods

### Meta-analysis

For this study we first re-performed a genome-wide meta-analysis of CL/P, CP, and all OFCs, from European and Asian ancestries^23^. The details of the sample collection and genotype quality control (QC) have been described previously^24^. Briefly, the original study recruited individuals with OFCs, their unaffected relatives, and unrelated controls (individuals with no known family history of OFCs or other craniofacial anomalies; N = 1626) from 18 sites across 13 countries from North America, Central or South America, Asia, Europe and Africa. Samples were genotyped for approximately 580,000 single nucleotide polymorphic (SNP) markers from the Illumina HumanCore+Exome array, of which approximately 539,000 SNPs passed quality control filters recommended by the Center for Inherited Disease Research (CIDR) and the Genetics Coordinating Center (GCC) at the University of Washington^24^. These data were then phased with SHAPEIT2 ^91^ and imputed with IMPUTE2^92^ to the 1000 Genomes Project Phase 3 release (September 2014) reference panel. The most-likely imputed genotypes were selected for statistical analysis if the highest probability (r2) > 0.9. SNP markers showing deviation from Hardy-Weinberg equilibrium in European controls, a minor allele frequency or MAF < 5%, or imputation INFO scores < 0.5 were filtered out of all subsequent analyses. The information for the genotyped markers was retained after imputation and the imputed values for these variants were only used to assess concordance.

### SNPs Selection of GWAS-identified loci/variants

We then selected 887 SNPs from eight loci that include a gene known to be expressed in oral epithelium and in most cases, regulates differentiation of an epithelial tissue (1q32/*IRF6*, 2p21/*THADA*, 6p24.3/*TFAP2A*, 9q22.2/*GADD45G*, 12q13.13/*KRT18*, 20q12/*MAFB* and 9q22.33/*FOXE1*) (Table 1, Supplementary Tables 1, 2). We selected SNPs with *P* values suggestive of association from seven loci (*IRF6*, *THADA*, *TFAP2A*, *GADD45G*, *KRT18*, and *MAFB*) from the meta-analysis^23^ (Table 1, Supplementary Table 2). We additionally included 59 SNPs at the *TFAP2A* locus identified in an independent GWAS of CL/P in Han Chinese^26^. Because of constraints on the library size, for one locus (*FOXE1*) we instead picked nine SNPs in strong linkage disequilibrium with the lead SNP (rs12347191) in a European population and annotated by Haploreg as being within regulatory elements^23^ (Supplementary Table 2).

### Reporter Library Construction

We designed an oligonucleotide library in which each oligo contained the sequence of the SNP of interest, a unique barcode, and priming and restriction sites necessary for amplification and cloning into a pGL-hsp68 reporter plasmid^93^. The sequence of human genomic DNA corresponding to 161 bp windows, centered on each of 887 GWAS-identified SNPs, was downloaded from the UCSC Table browser tool (genome build: hg19). We generated two versions of each cis regulatory element (CRE) that were identical except for harboring the major or minor allele of the SNP, respectively (1774 elements) (Fig.1a, Supplementary Table 3). We ordered a pool of 7484 unique 230-mer oligonucleotides (elements) through a limited licensing agreement with Agilent Technologies (Santa Clara, CA). To establish an empirical null distribution of activity in the assay, we included 90 sequences that were scrambled versions of the sequences of other elements (Supplementary Table 4). To account for noise in the assay, each sequence was assigned four unique barcodes. Finally, to create constructs with only the basal promoter and no upstream regulatory sequence, we included an additional random 28 filler elements, which were excised in the second step of cloning, yielding 28 basal promoters only controls with one barcode each. In total we synthesized 7484 elements (7096+360+28) (Supplementary Table 4, Fig. 1a).

Oligos were designed with the following structure (Supplementary Fig. 21): a forward priming sequence (PS1) with NheI restriction site, (primer for this site is KC_1F), 161 base cis-regulatory element (Supplementary Table 3), AgeI restriction site, NruI restriction site, MluI restriction site, 9 base barcode, and a reverse priming sequence (PS2) with XhoI site (primer for this site is KC_1R) (Fig. 1a, Supplementary Fig. 21, Supplementary Table 4).

We amplified the oligo pool using five cycles of PCR with Phusion High-Fidelity polymerase (New England Biolabs) and primer set KC_1 (Supplementary Table 18). We cloned the pool of amplicons into the pGL-hsp68 backbone using NheI and XhoI which excludes the hsp68 promoter fused to DsRed. We prepared DNA from 41,000 colonies to generate library KC_L1. We then amplified a DNA fragment containing the hsp68 promoter and the DsRed cDNA using primers KC_2F and KC_2R (Supplementary Table 18) containing AgeI and MluI restriction sites, respectively, and cloned this fragment into the KC_L1 library such that the CREs lie upstream to the hsp68 promoter driving DsRed and the barcodes; this procedure created library KC_L2.

To assess the completeness of the cloned library, we sequenced the barcodes in the final plasmid pool in library KC_L2. It was achieved by amplifying the barcode fragments using primer set KC_BC containing with NheI and MfeI restriction sites (Supplementary Table 18), followed by specific Illumina adapters ligation (P1 and PE2 adapters; Supplementary Table 18), and Illumina enrichment PCR using KC_3 set of primers (Supplementary Table 18), and sequencing.

### Cell lines

The human fetal oral epithelial cell line (GMSM-K)^61^ (a kind gift from Dr. Daniel Grenier, Université Laval) was maintained in keratinocyte serum-free medium (KSFM) supplemented with human recombinant epidermal growth factor (EGF 30 µg/ml) and bovine pituitary extract (BPE 0.2 ng/ml) (Thermo Fisher Scientific). Human Epidermal Keratinocytes from neonatal foreskin (HEKn; primary neonatal keratinocytes) purchased from ATCC (ATCC Catalog no. PCS-200-010) were maintained and cultured as per manufacturer’s instruction in dermal cell basal medium (ATCC Catalog no. PCS-200-030) supplemented with keratinocyte growth kit (ATCC Catalog no. PCS-200-040). The Normal Human Skin Keratinocyte (NHSK)^94^ cell line was grown in KSFM or on an irradiated NIH3T3 feeder layer in Dulbecco’s Modified Eagle’s Medium (DMEM) and HamF12 (3:1 ratio) according to Rheinwald and Green^95^. WTC-11 human induced pluripotent stem cells (iPSCs) (Coriell Institute for Medical Research, Catalog no. GM25256), were maintained in mTeSR Plus media (StemCell Technologies) on Matrigel (Corning)-coated plates and passaged with Accutase. Cells were incubated at 37°C in 5% CO_2_. Details of cell lines/primary cells/induced pluripotent cells used in the study are provided in Supplementary Table 5.

### Transfection of CRE-library (plasmid pool)

A pool of plasmids (from library KC_L2; as described in Methods, Reporter library construction) in four replicates, 2.5 µg each, was transfected into 1 million GMSM-K with Amaxa Cell Line Nucleofector Kit V (Lonza, Cologne, Germany) using Nucleofector II (Lonza) (program: T-020). Forty eight hours post transfection RNA and genomic DNA was extracted from each set using Quick-DNA/RNA Miniprep kit (Zymo Research Catalog no. D7001) as per manufacturer’s instructions.

### Sample preparation for RNA-Seq

Isolated RNA was treated with Turbo DNaseI (ThermoFisher Catalog no. AM1907) as per manufacturer’s instructions. First strand of cDNA was synthesized with SuperScript III First-Strand Synthesis System (Thermo Fisher Scientific Catalog no. 18080051). The barcodes including the flanking sequence were amplified from both cDNA, genomic DNA (transfected plasmid DNA) and plasmid DNA samples (that was used for transfection) with Phusion High-Fidelity PCR Master Mix (NEB Catalog no. M0531S) using primer set KC_BC (Supplementary Table 18) and that yielded barcode sequences that were flanked with NheI and MfeI restriction enzyme sites. The products were digested with NheI and MfeI and then ligated with Illumina adapter sequences P1 and PE2 (Supplementary Table 18). This product was further amplified using primer set KC_3 (Supplementary Table 18) followed by sequencing.

To measure expression of the barcoded library, electroporation replicates were multiplexed and run on two lanes of an Illumina HiSeq machine, which generated 48.2 million sequence reads corresponding to cDNA and 48.8 million reads corresponding to DNA (from plasmids). Sequencing reads that matched the first 20 nucleotides of designed sequence were counted, regardless of quality score.

### Statistical analysis to determine regulatory effects of SNPs by MPRA

We filtered out barcodes that were low abundance in the plasmid pool (<50 reads), then set any cDNA barcodes that were detection-limited in at least one sample (<50 reads for replicates run on the first lane, <100 reads for the replicates run on the second lane) to zero across all cDNA samples. We normalized each sample to the total counts per million (CPM). RNA barcode levels were highly correlated among 4 replicates (Supplementary Fig. 22). There was a strong correlation in the barcode counts in the plasmid library before transfection and after harvesting plasmid from transfected cells (Supplementary Fig. 23). Therefore, we used plasmid barcode counts (pre-transfection) for further analyses. We further normalized RNA barcode counts by the plasmid barcode counts (RNA/DNA). We checked the reproducibility based on log2RNA/DNA counts (Supplementary Fig. 24). To calculate the reporter activity for each sequence we averaged the log2RNA/DNA counts within a replicate, normalized to the within-replicate mean basal barcode activity and then averaged across all the replicates (Supplementary Fig. 1a). The standard error of the mean (SEM) was calculated using the basal-normalized activities across replicates. Variant significance was determined by performing Welch’s t-test between major and minor alleles followed by FDR correction (Supplementary Table 6). We used our previously published^96^ Python code for processing and analyzing MPRA reads, which is available on Github at [https://github.com/barakcohenlab/CRX-Information-Content].

### Activity-by-Contact to assign SNP-containing-enhancers to promoters

The activity-by-contact method (ABC) utilizes cell-type specific chromatin accessibility (ATAC-seq or DNase hypersensitivity) and chromatin activity (H3K27ac) and HiC data that is either cell-type specific or averaged over 10 cell types – the performance of this tool is similar with either type of HiC data^63^. To calculate the ABC scores for keratinocyte enhancers we used publicly available datasets from primary Normal Human Epidermal Keratinocytes (NHEK) cells for H3K27ac (GSM733674), DNase hypersensitivity (GSM736545), RNA-seq (GSM958736 for) and averaged HiC data from 10 cell types as reported in Fulco et al^63^. All elements with an ABC score >0.01 were included in the analysis.

### Electroporation and dual luciferase reporter assay

For dual luciferase reporter assays, each reporter construct (Supplementary Table 10) was cloned into the pGL3 promoter vector and constructs were co-transfected with Renilla luciferase plasmid. Briefly, GMSM-K or HEKn cells were electroporated with plasmid using the Amaxa Cell Line Nucleofector Kit (Lonza) and the Nucleofector II instrument (Lonza) program T-020. We used a dual-luciferase reporter assay system (Promega) and FB12 Luminometer (Berthold Detection Systems) to evaluate the luciferase activity following manufacturer’s instructions. Relative luciferase activity was the ratio of Firefly and Renilla activities. At least three independent measurements were performed for each transfection group. All results were presented as mean ± standard deviation (SD). Statistical significance was determined using two-tailed Student’s *t*-test.

### Immunofluorescence detection and imaging

GMSM-K and Normal Human Skin Keratinocytes (NHSK) were grown on glass coverslips, fixed in 4% paraformaldehyde for 10 min, permeabilized in 0.2% Triton-X100 for 2 min, then rinsed in PBS. Cells were incubated in 3% goat serum (Vector Laboratories, Catalog no. S-1000-20) in 2% PBS-Bovine Serum Albumin for 30 min to block non-specific binding, followed by incubation in primary then secondary fluorescent antibodies 1 h each at room temperature with PBS washes in between the two incubations. Primary antibodies that were used are: mouse monoclonal anti-E-cadherin (BD Bioscience, Catalog no. 610181, 1/100), mouse monoclonal anti P63 (clone 4A4, Santa Cruz, Catalog no. SC-8431, 1/250), mouse monoclonal anti human Keratin 14 (clone LL002, Novocastra, Catalog no. NCL-LL002, 1/200). Goat anti-mouse Alexa Fluor 488 (Invitrogen, Catalog no. A-11001) was used as secondary antibody. Following PBS washes, coverslips were dried and mounted on a microscope slide using ProLong Diamond antifade mounting medium containing DAPI (ThermoFisher Scientific, Catalog no. P36962) and cured for 24 h at room temperature in the dark. Images were collected with Zeiss 880 or 980 confocal microscopes using the ZEN software (Zeiss). Confocal images were processed using Fiji software.

### CRISPR-Cas9 mediated genome editing

Genome engineering for each SNP (rs11119348, rs661849 and rs10984103) in iPSCs or GMSM-K was achieved using a previously described protocol^89^. Sequences of CRISPR RNAs (crRNA) and repair template specific to each SNPs are provided in (Supplementary Table 19). Briefly, equimolar concentrations of specific crRNA and trans-activating crRNA (tracrRNA, IDT Catalog no. 1072532) were annealed at 95°C for 5 min followed by cooling at room temperature (RT) to form functional gRNA duplexes. Further, the ribonucleoprotein (RNP) complex was prepared 15 min before transfection by mixing gRNAs (1.5 µM) with Cas9 protein (0.45 µM) (Sigma, Catalog no. CAS9PROT for iPSCs and IDT Catalog no. 1081058 for GMSM-K) at RT. RNP complexes together with the repair template were transfected into 1 million iPSCs with Amaxa nucleofector (Lonza, Human Stem Cell kit 2, program: A-023 for iPSCs) in presence of ROCK inhibitor (Y-27632, Catalog no. 04-0012-10, Stemgent; 10 mM stock). Lonza Kit V, program: T-020 was used for transfecting GMSM-K. Individual colonies were hand-picked and plated into 96 well plates. DNA was extracted using Quick Extract DNA extraction solution (Epicentre Catalog no. QE09050). Colonies were screened by PCR and sequenced using primers flanking each SNP (Primer sequences are provided in Supplementary Table 20a). From iPSCs transfection experiment, two independent clones each of homozygous risk (AA for rs11119348; CC for rs661849) and homozygous non-risk (CC for rs11119348; TT for rs661849) for each SNP near *IRF6*, and three independent clones of each of homozygous risk (CC) and non-risk (AA) genotype for rs10984103 near *FOXE1* were isolated (Supplementary Figs. 13a, b, 16a). From GMSM-K transfection experiment, multiple independent clones, six clones of homozygous risk (AA for rs11119348), seven clones of homozygous non-risk (CC for rs11119348) and four independent clones each of the homozygous risk (CC for rs661849), and homozygous non-risk (TT for rs661849) were isolated. Deletion of *FOXE1* gene in GMSM-K was achieved using a pair of gRNA (Supplementary Fig. 19). A set of three primers was used for colony screening and three independent clones were isolated (Supplementary Table 20b). Sequencing of three independent clones revealed that all but 30 bp at the 3’ end of the single coding exon gene was deleted (Supplementary Fig. 19).

### *In vitro* differentiation of iPSCs into embryonic oral epithelial cells

Two independent samples of each genotype (2 clones of the homozygous risk genotype, and 2 clones of the homozygous non-risk genotype) for both SNPs at *IRF6* locus and three independent samples of each genotype (3 clones of the homozygous risk genotype, and 3 clones of the homozygous non-risk genotype) for rs10984103 at *FOXE1* locus were seeded in triplicate in 12-well plate (50,000 cells per well). Additionally, three independent samples of heterozygous genotype for each SNP individually were seeded (Supplementary Figs. 13a, b, 16a). Further, the cells were induced to differentiate into embryonic oral epithelial (iOE) cells using a 10-day differentiation protocol described previously^70, 89^ (Supplementary Fig. 13a, b, 16a).

### Quantitative RT-PCR on induced embryonic oral epithelial (iOE) cells or GMSM-K

RNA was extracted from multiple samples of each genotype (homozygous risk and homozygous non-risk) for rs11119348, rs661849 and rs10984103 (iOE cells – 6 each genotype for both SNPs near *IRF6* and 9 each genotype for rs10984103 near *FOXE1*; GMSM-K - 6 non-risk and 7 risk genotype for rs11119348 and 4 each genotype for rs661849) using Quick-DNA/RNA Miniprep kit (Zymo Research #D7001) followed by DNase treatment using manufacturer’s instructions (Thermo Fisher Scientific #AM1907). Reverse transcription was performed using High-Capacity cDNA Reverse Transcription Kit (Thermo Fisher Scientific # 4368814). Real-time PCR was performed using SYBR Green qPCR mix (Bio-Rad) in Bio-Rad CFX Connect Real-Time System. Quantitative RT-PCR reaction for each sample was performed in triplicate. Data analysis was performed using previously described methods^97, 98^. Briefly, expression levels of *IRF6* were normalized against *ACTB* ^97^ for GMSM-K and expression levels of *IRF6* or *FOXE1* were normalized against *ACTB, GAPDH, HPRT, UBC* and *CDH1*^98^ for iOE cells (mentioned in the respective figure legends). Data were presented as mean ± SD. Statistical significance was determined using two-tailed Student’s *t*-test. Primer sequences used in the study for qRT-PCR are provided in Supplementary Table 20c.

### Transcription factor binding site analysis using JASPAR 2022

Ten nucleotides flaking each side of the *IRF6* -10 kb, *IRF6* -22 kb and *FOXE1* 24 kb SNPs (21 nucleotides long), individually with non-risk and risk allele was used for transcription factor binding site (TFBS) analysis using JASPAR 2022^71^. An 80% threshold was used to detect transcription factor binding sites, and all those with a score below 4.0 were discarded, to focus on binding sites of at least medium affinity. For TFBS that, at 80% threshold, were specific for the risk or non-risk allele, we dropped the threshold to 50% to learn the affinity of such TFBSs with the other allele.

### Chromatin immunoprecipitation qPCR

Induced oral epithelial cells (iOE cells) derived from iPSC that are heterozygous for risk and non-risk alleles of rs11119348, rs661849, and rs10984103 or GMSM-K homozygous for risk or non-risk alleles for rs11119348 and rs661849 were harvested (3.5 million cells) and fixed in 1% formaldehyde for 15 min at RT. Fixation was stopped with 125 mM glycine for 5 min at RT. Chromatin immunoprecipitation (ChIP) on iOE cells was performed using a semi-automated protocol described previously^99^. Briefly, cells were sheared using PIXUL or VirSonic 600 (5 min with 30/40 second on/off cycles for GMSM-K for *IRF6* -22 kb SNP and iOE cells for *FOXE1* 24 kb SNP) to obtain genomic DNA fragments averaging 300 to 600 bp. The sonicated cell lysates were subjected to chromatin immunoprecipitation with specific antibody (1 μg of anti-H3K27Ac from Millipore-Sigma Catalog no. 07-360 and 1 μg of anti-ETS2 from GeneTex Catalog no. GTX104527 and anti-FOXE1, a kind gift from Pilar Santisteban, Universidad Autónoma de Madrid, as previously described^100^). Normal rabbit IgG (Millipore-Sigma; Catalog no. 12-370) was used as negative antibody control. Quantitative PCR was performed with equal volume of DNA in triplicate from three ChIP replicates of each antibody and percent input DNA was calculated using the method described previously^101^. Primer sequences are provided in Supplementary Table 20d. Negative control PCR primers were designed to target a sequence 103.7 kb upstream of the *IRF6* transcription start site or 28.2 kb downstream to the *FOXE1* gene that did not harbor active regulatory elements identified from ATAC-Seq and H3K27Ac ChIP-Seq in Human immortalized oral epithelial cells (HIOEC) or NHEK. Data were presented as mean ± SD. Statistical significance was determined using two-tailed Student’s *t*-test.

### *ETS2* and *FOXE1* knockdown

siRNAs targeting *FOXE1* (Santa Cruz Catalog no. sc-44175) or *ETS2* (Santa Cruz Catalog no. sc-37855) and siRNA controls (Santa Cruz Catalog no.sc-37007) were transfected into one million HEKn (heterozygous for *IRF6* -22 kb SNP) or GMSM-K (homozygous risk for *IRF6* -10 kb or -22 kb SNPs). RNA extraction and qRT-PCR were performed on three replicates. Data analysis was performed using previously described method^97^. Briefly, expression levels of genes of interest (*FOXE1*, *ETS2*, or *IRF6*) were normalized against *ACTB*^97^ for GMSM-K or HEKn. Data were presented as mean ± SD. Statistical significance was determined using two-tailed Student’s *t*-test. Primer sequences used in the study for qRT-PCR are provided in Supplementary Table 20c. RT-PCR primer pair B from Santa Cruz (Catalog no. sc-44175-PR) for *FOXE1* was used.

Further, siRNA targeting *FOXE1* or *ETS2* and control siRNA were co-transfected with respective risk harboring luciferase constructs along with Renilla luciferase plasmid. Three independent transfections were performed. Cells were harvested 48 hours post transfection and dual luciferase reporter assays were performed as described earlier. The RNA expression and luciferase reporter assays were carried out on *FOXE1* KO GMSM-K as well. GraphPad Prism 10 was used for the statistical data analysis for luciferase reporter assays, RNA expression and ChIP-qPCR experiments.

### Conditional Analysis Methods

Conditional analyses were conducted using genome-wide genotyping data from the Pittsburgh Orofacial Cleft (POFC) Study on the case-control subset of the dataset (described earlier). A total of 394 cleft lip only cases and 1995 cleft lip and palate cases were used in the analysis with 1626 unrelated controls. Common variants (MAF > 5%) near the *IRF6* locus were included in this analysis and the association between each variant and OFCs was calculated using a logistic regression in PLINK (v1.9)^102^ controlled for sex and 10 PCs of ancestry. Conditional analyses also controlled for either the genotype of rs661849, rs11119348, or the genotypes of both rs661849 and rs11119348. Since the data came from a genome-wide analysis, SNPs with association *P* values less than 5 × 10^−8^ were considered genome-wide significant. Regional association plots were made with LocusZoom, where the linkage disequilibrium (LD) blocks and recombination rates were estimated from European populations^103^. LD between pairs of SNPs was calculated in the data from the 1000 Genomes Project using LDLink^104, 105^.

## Supplemental information

Twenty supplementary tables provided as Excel sheets and supplementary Figs. (1-24) provided in one PDF document.

## Data availability

MPRA and RNA-Seq data of GMSM-K are provided in Supplementary Table 6 and 12 respectively. Data used for conditional analysis is publicly available: dbGaP Study Accession: phs000774.v2.p1.

## Author Contributions

P.K. contributed to the design, data acquisition, analysis, and interpretation, drafting and editing the manuscript; R.Z.F. - data analysis, interpretation and editing the manuscript, L.P. - data analysis; S.W.C. - data analysis, interpretation, editing the manuscript; K.P., A.V. - generation and preliminary analyses of murine *in vivo* reporter assays for *IRF6* -10 kb and -22 kb SNPs; L.R. - data acquisition, analysis; M.D. - data acquisition, analysis, interpretation and editing the manuscript; A.P.P - data acquisition; D.M., K.B. - design and data acquisition; J.M. - data acquisition and analysis; H.R.B. - design, and data interpretation; E.J.L. - design and data interpretation; M.A.W. - design, data acquisition, analysis, and interpretation, editing the manuscript, B.A.C. - design, data interpretation, R.A.C. - conception, design, data acquisition, analysis, interpretation, drafting and editing the manuscript. All authors gave final approval and agree to be accountable for all aspects of the work.

## Acknowledgments

The authors thank Pilar Santisteban, Universidad Autónoma de Madrid, for the FOXE1 antibody, members of the Craniofacial Interest Group at the University of Iowa for helpful comments, and Greg Bonde (University of Iowa) and Josh Rosswork (University of Washington) for technical assistance throughout the project. We thank Ellison Stem Cell Core at ISCRM and Jennifer Hesson for technical assistance in screening and genotyping of iPSCs. We are grateful for many discussions with members of the Craniofacial Interest Group weekly meeting hosted at the University of Iowa.

This project was funded by grants from the U.S. National Institutes of Health, DE027362 (R.A.C.), R01DE028599 (A. V.), R01DE033016 (R.A.C., H.R.B., J.M.), R01AR067739 (M.D.), HG010855 (K.B.) and CA246503 (K.B.). Research conducted at the E.O. Lawrence Berkeley National Laboratory was performed under Department of Energy Contract DE-AC02-05CH11231, University of California.

