## Supplementary figures for "Identification of functional non-coding variants associated with orofacial cleft"

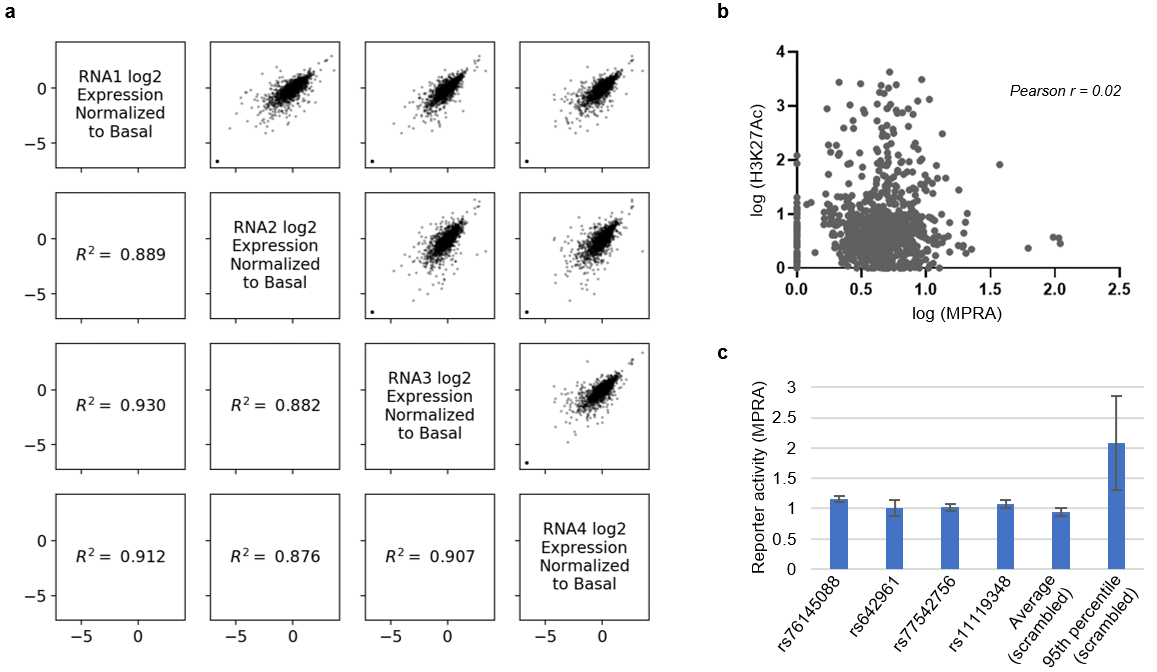

**Supplementary Fig. 1: a)** Correlation plot showing reporter activity of individual SNP-containing elements, normalized to basal expression, between replicates of the MPRA. **b)** Correlation plot between MPRA data from GMSM-K and H3K27Ac signal (GSM733674) from normal human epidermal keratinocytes (NHEK). **c)** Bar chart showing average reporter activity in the MPRA for the four indicated elements, all from within *IRF6* MCS9.7. Fifth bar, average reporter activity from 84 scrambled elements. Sixth bar, average reporter activity of the scrambled element with 95^th^ percentile activity. Data are represented as mean ± standard error of the mean (SEM) from four replicate experiments and four barcodes per element.

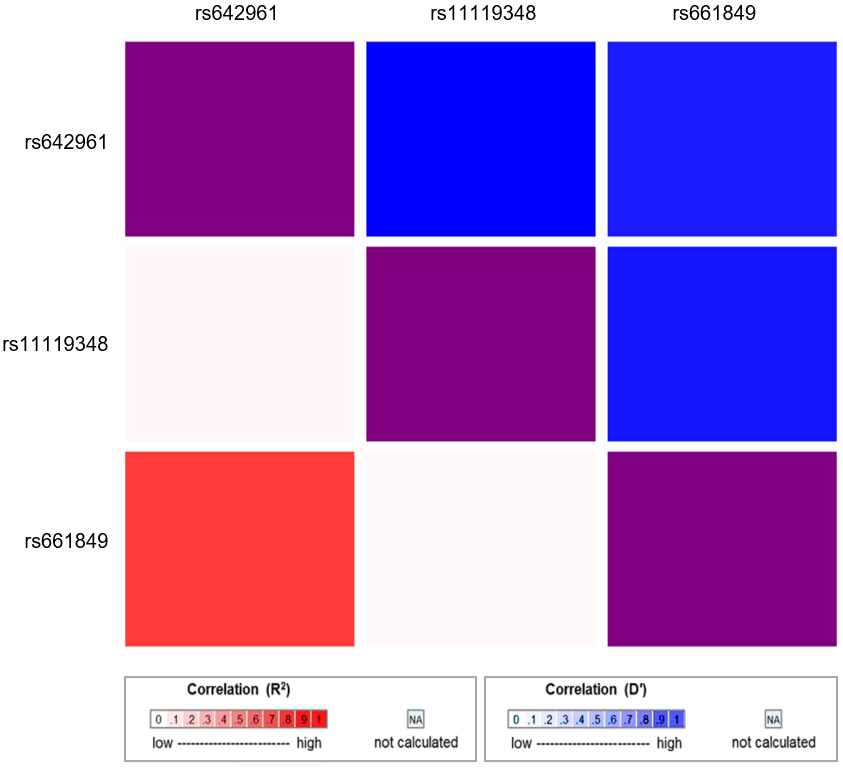

**Supplementary Fig. 2:** Linkage disequilibrium plot between rs642961 and *IRF6* -10 kb (rs11119348) and *IRF6* -22 kb (rs661849) SNPs. R^2^ is the correlation between a pair of loci. D’ is the difference between the observed and the expected frequency of a given haplotype. Although rs642961 is in *IRF6* MCS9.7, like rs11119348, it is in strong linkage disequilibrium with rs661849 but not with rs11119348.

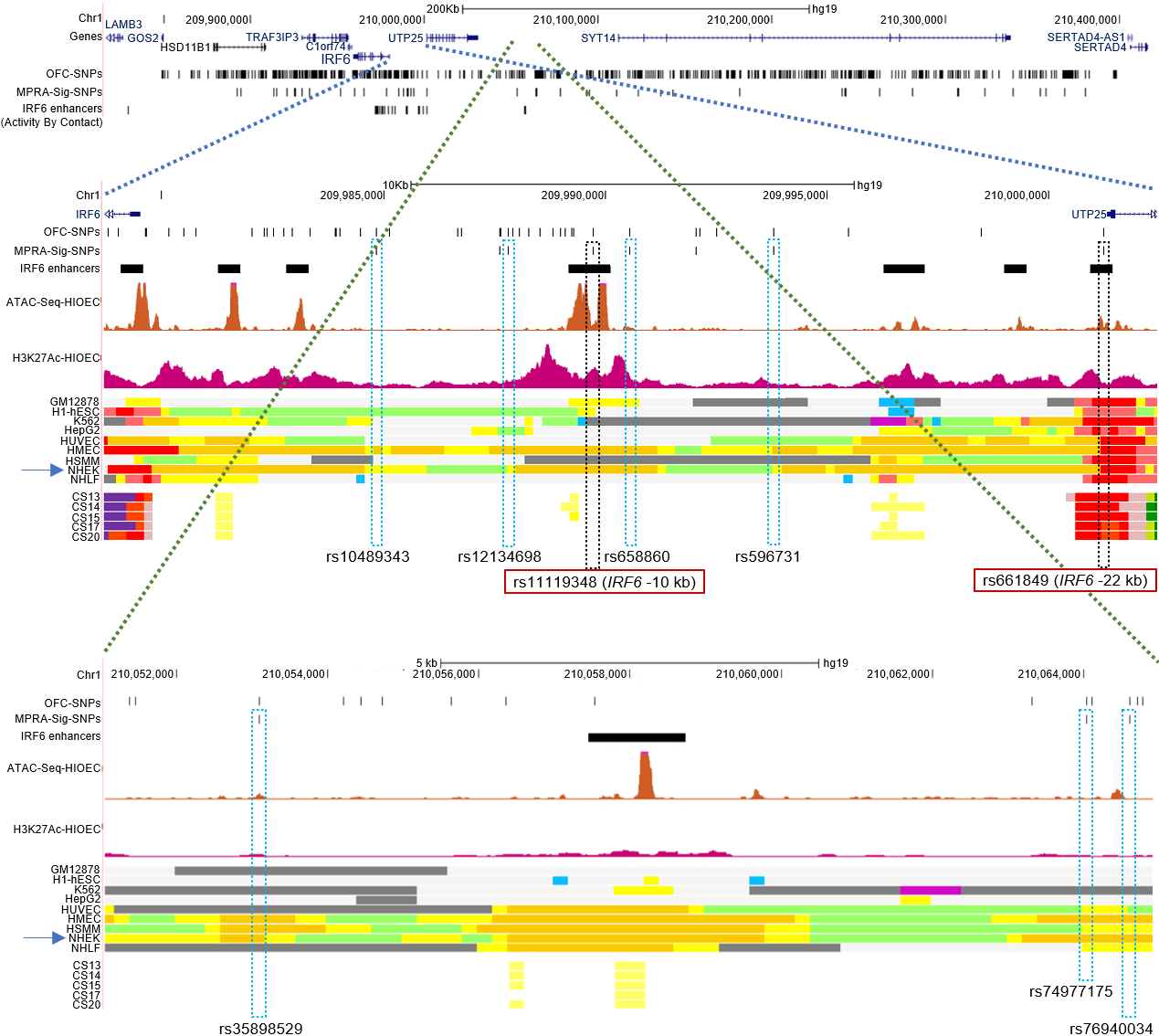

**Supplementary Fig. 3:** **SNPs near *IRF6* with significant effect in MPRA and overlying enhancers active in NHEK**. Browser view of the human genome, GRCh37/hg19, focused on the GWAS-identified SNPs at *IRF6* associated with orofacial cleft. Tracks and color codes are as described in Fig. 2. Blue and green dashed lines indicate the zoomed-in browser views from the left-hand side and right-hand side OFC-associated SNPs at *IRF6* respectively. Blue boxes, SNPs with significant effect in MPRA lying within chromatin marked as an active enhancer (yellow or gold) in NHEK (blue arrow).

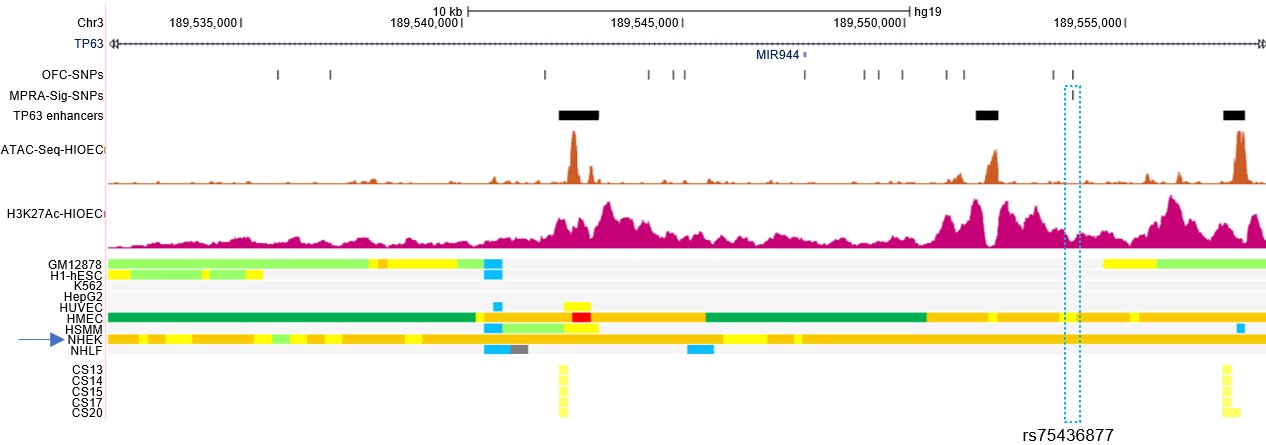

**Supplementary Fig. 4:** **SNPs near *TP63* with significant effect in MPRA and overlying enhancers active in NHEK.** Browser view of the human genome, GRCh37/hg19, focused on the GWAS-identified SNPs at *TP63* associated with orofacial cleft. Tracks and color codes are as described in Fig. 2. Blue box, a SNP with significant effect in MPRA lying within chromatin marked as an active enhancer in NHEK (blue arrow).

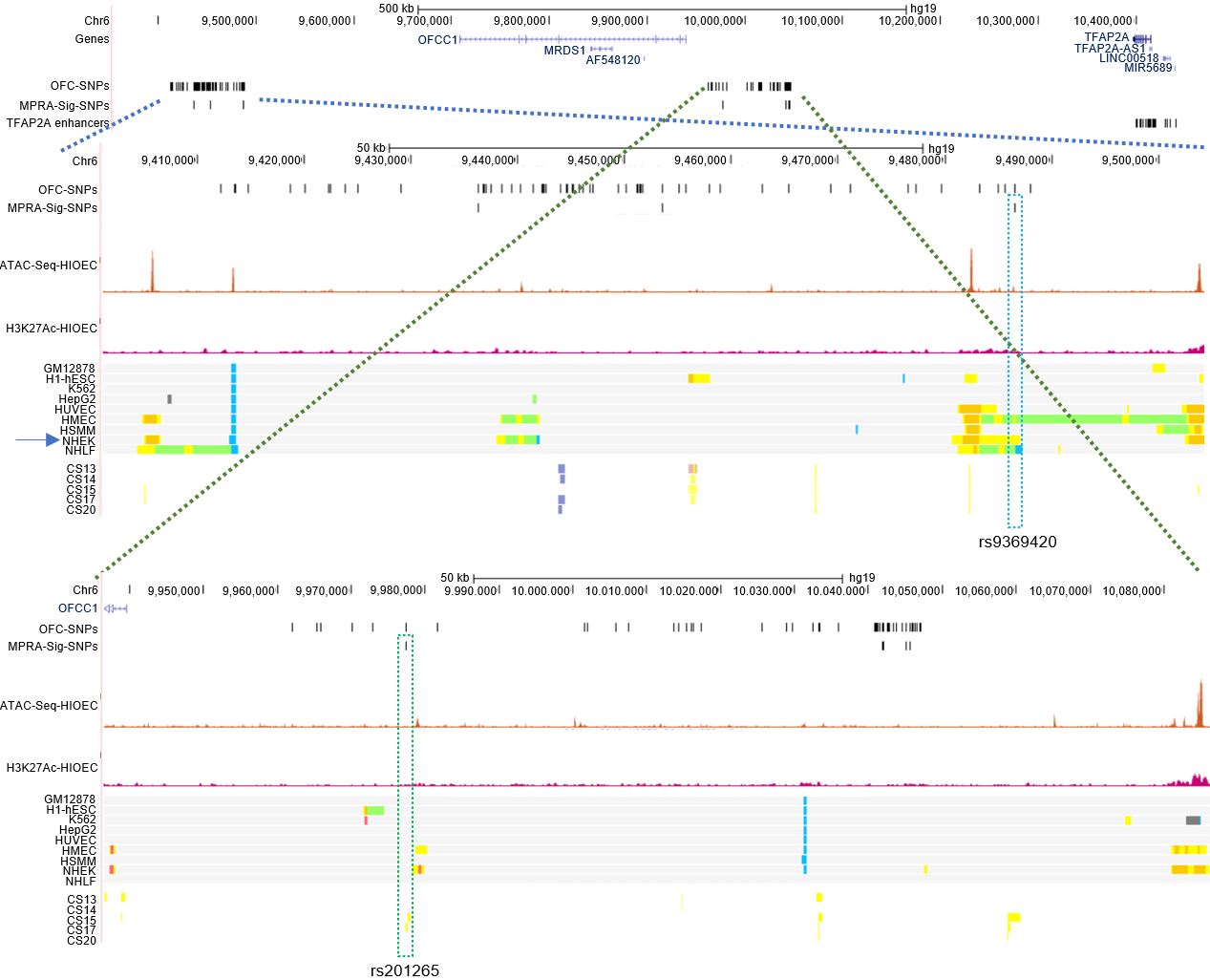

**Supplementary Fig. 5:** **SNPs near *TFAP2A* with significant effect in MPRA and overlying enhancers active in NHEK or human embryonic faces.** Browser view of the human genome, GRCh37/hg19, focused on the GWAS-identified SNPs at *TFAP2A* associated with orofacial cleft. Tracks and color codes are as described in Fig. 2. Blue and green dashed lines indicate the zoomed-in browser views from the left-hand side and right-hand side OFC-associated SNPs at *TFAP2A,* respectively. Boxes, SNPs with significant effect in MPRA lying within chromatin marked as an active enhancer in NHEK (blue arrow and blue box) or human embryonic faces (green box).

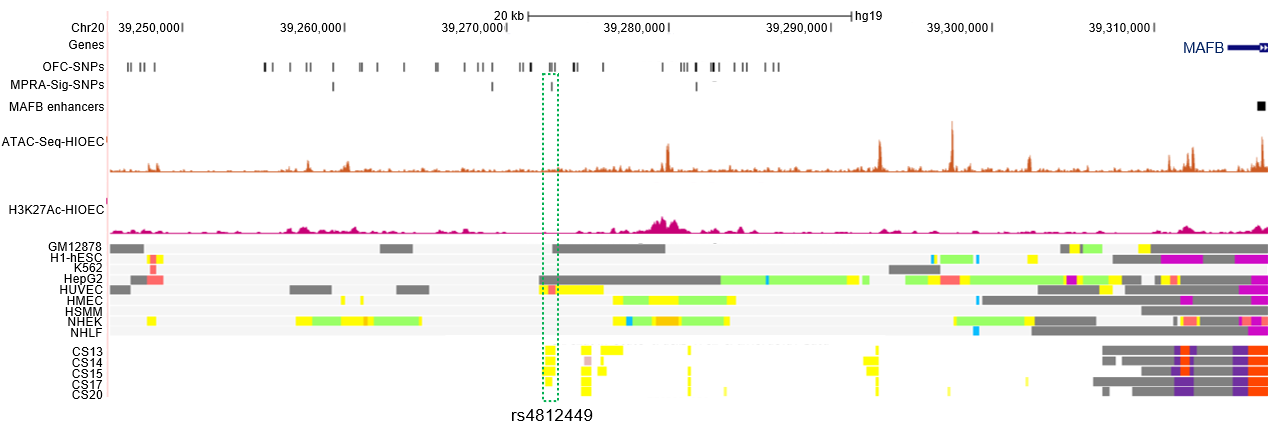

**Supplementary Fig. 6:** **SNPs near *MAFB* with significant effect in MPRA and overlying enhancers active in human embryonic faces.** Browser view of the human genome, GRCh37/hg19, focused on the GWAS-identified SNPs at *MAFB* associated with orofacial cleft. Tracks and color code are as described in Fig. 2. Green box, a SNP with significant effect in MPRA lying within chromatin marked as an active enhancer in human embryonic faces.

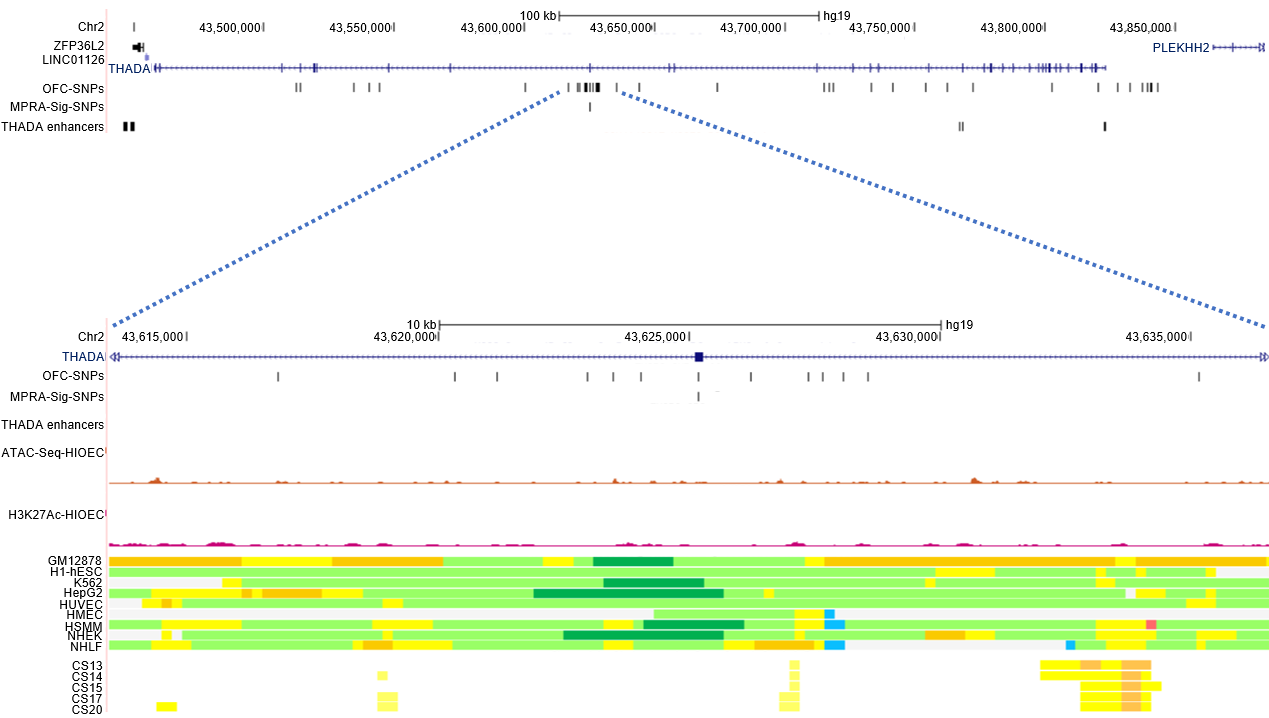

**Supplementary Fig. 7:** **SNPs near *THADA* with significant effect in MPRA.** Browser view of the human genome, GRCh37/hg19, focused on the GWAS-identified SNPs at *THADA* associated with orofacial cleft. Tracks and color codes are as described in Fig. 2. Blue dashed line indicates the zoomed-in browser view of OFC-associated SNPs at *THADA.* In this locus, the SNP with allele-specific effect in the MPRA does not lie in chromatin marked as an enhancer in NHEK or in human embryonic faces.

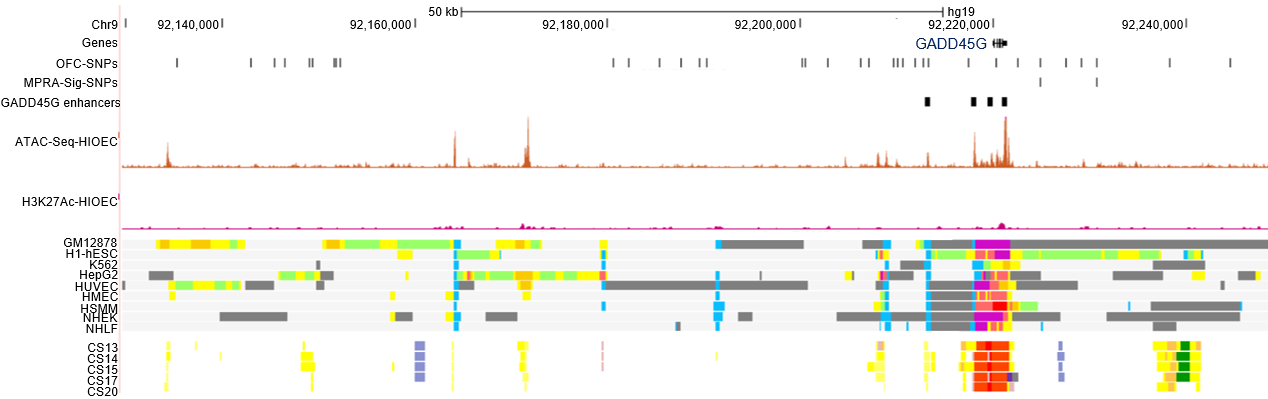

**Supplementary Fig. 8:** **SNPs near** ***GADD45G* with significant effect in MPRA.** Browser view of the human genome, GRCh37/hg19, focused on the GWAS-identified SNPs at *GADD45G* associated with orofacial cleft. Tracks and color codes are as described in Fig. 2. In this locus, the SNPs with allele-specific effects in the MPRA do not lie in chromatin marked as an enhancer in NHEK or in human embryonic faces.

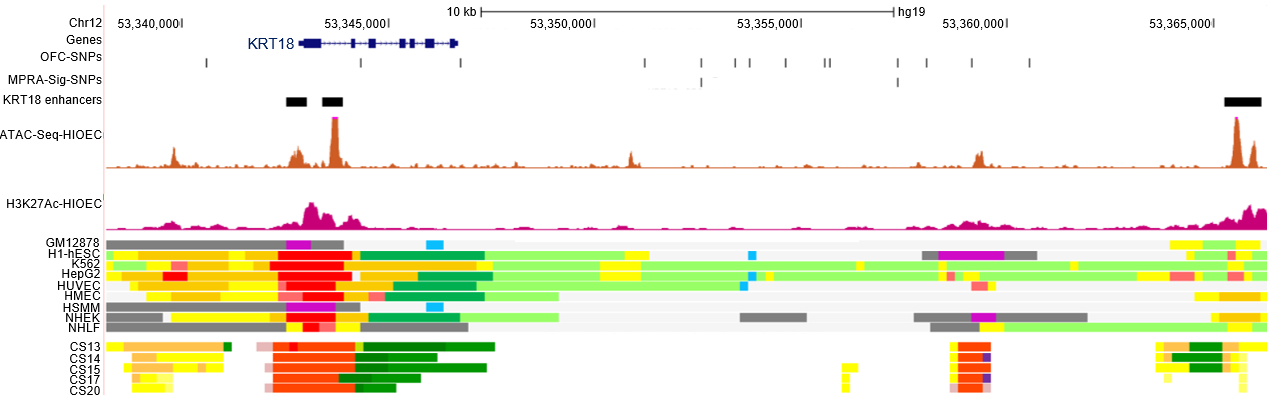

**Supplementary Fig. 9:** **SNPs near *KRT18* with significant effect in MPRA.** Browser view of the human genome, GRCh37/hg19, focused on the GWAS-identified SNPs at *KRT18* associated with orofacial cleft. Tracks and color codes are as described in Fig. 2. In this locus, the SNPs with allele-specific effects in the MPRA do not lie in chromatin marked as an enhancer in NHEK or in human embryonic faces.

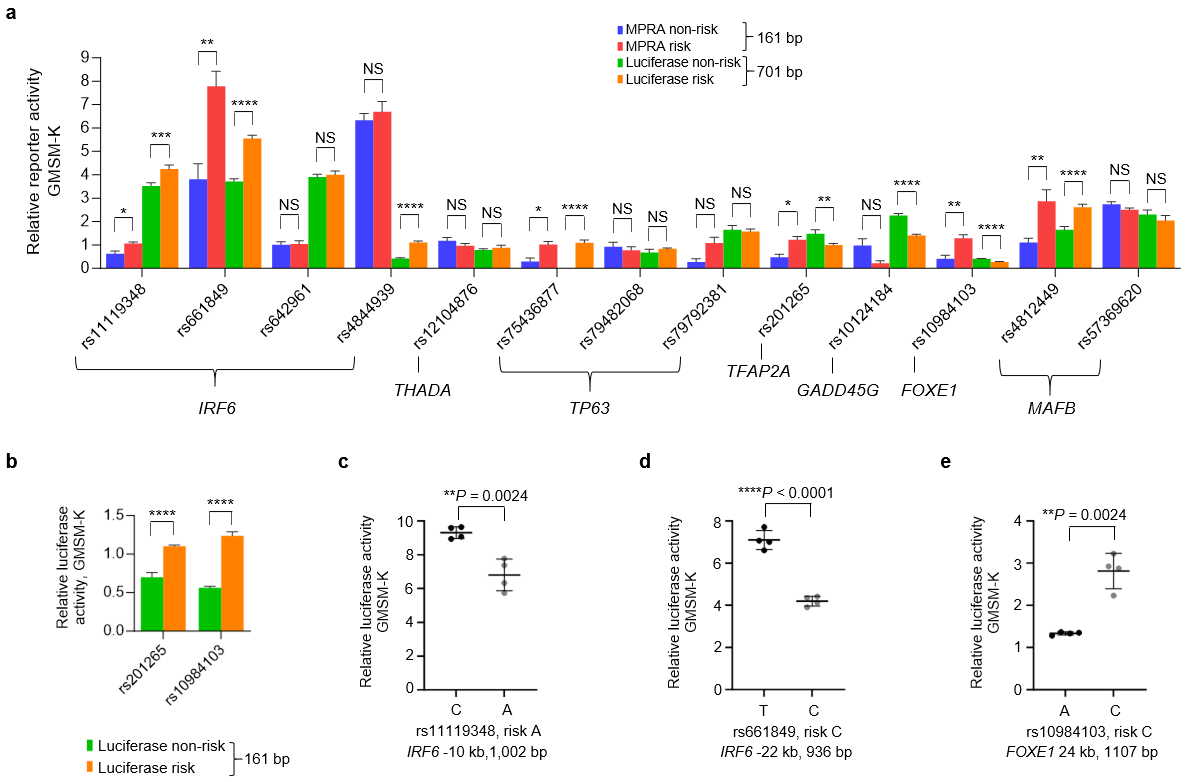

**Supplementary Fig. 10:** Luciferase reporter assay in GMSM-K: **a)** Bar chart of MPRA and luciferase reporter activities for the indicated SNPs at various loci in GMSM-K using elements of the indicated length centered on the SNP. For luciferase reporter activities, data are represented as mean ± standard deviation (SD) from four independent experiments. Statistical significance (*P* value, two-tailed) is determined by Student’s *t*-test (**P*<0.05, ***P*<0.01, ****P*<0.001, *****P*<0.0001). NS, non-significant. **b)** Bar chart of luciferase reporter activities using 161 bp elements for two SNPs (one each at *TFAP2A* and *FOXE1*) in GMSM-K. Data are represented as mean ± SD from four independent experiments. Statistical significance (*P* value, two-tailed) is determined by Student’s *t*-test (*****P*<0.0001). **c-e)** Scattered dot plot of relative luciferase activity using elements of the indicated length and designed to match open chromatin flanking the SNP in HIOEC for non-risk and risk alleles of rs11119348, rs661849 and rs10984103 respectively in GMSM-K. Data are represented as mean ± SD from four independent experiments. Statistical significance (*P* value, two-tailed) is determined by Student’s *t*-test.

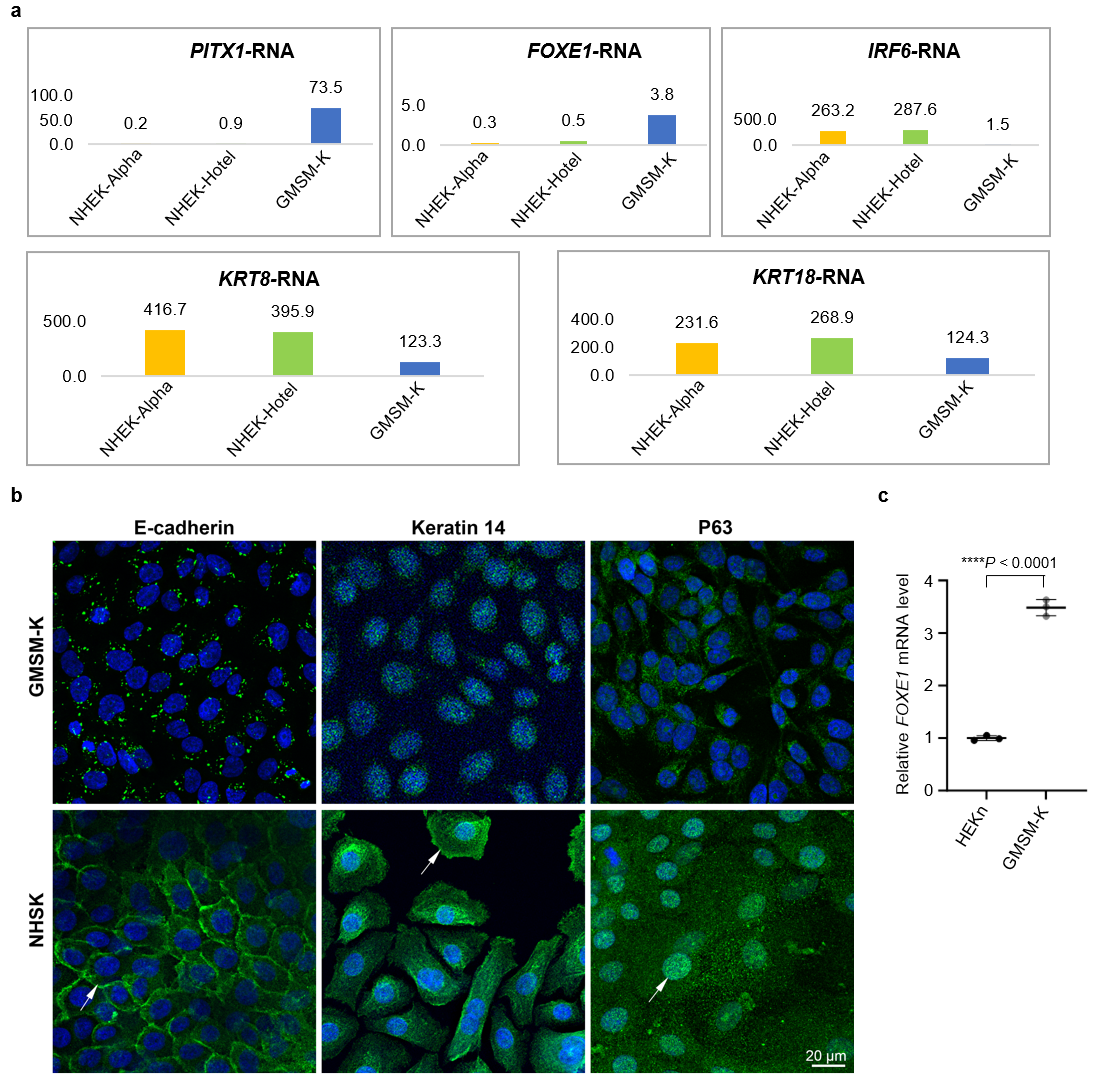

**Supplementary Fig. 11: a)** Comparative expression levels of the indicated genes from RNA-Seq datasets: GMSM-K, from the present study; NHEK-Alpha, NCBI GEO identifier GSM6050574; NHEK-Hotel, NCBI GEO identifier GSM6050576. **b)** Immunofluorescence staining for E-cadherin, Keratin 14, and P63 (green) of GMSM-K and normal human skin keratinocytes (NHSK)^1^. White arrows indicate positive staining in NHSK absent in GMSM-K. Nuclear DNA is labeled with DAPI (blue). Scale bar = 20 µm. **c)** Scattered dot plot of relative levels of *FOXE1* mRNA in HEKn and GMSM-K assessed by qRT-PCR. Expression levels of *FOXE1* are normalized against *ACTB.* Data are represented as mean ± SD from three replicates. Statistical significance (*P* value, two-tailed) is determined by Student’s *t*-test.

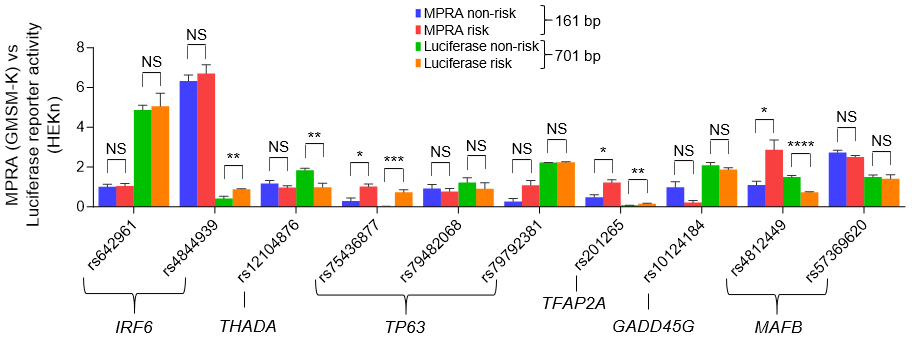

**Supplementary Fig. 12:** Bar chart of MPRA (GMSM-K) and luciferase reporter activities (HEKn) for the indicated SNPs at various loci. For luciferase reporter activities, data are represented as mean ± SD from three independent experiments. Statistical significance (*P* value, two-tailed) is determined by Student’s *t*-test. **P*<0.05, ***P*<0.01, ****P*<0.001, *****P*<0.0001. NS, non-significant.

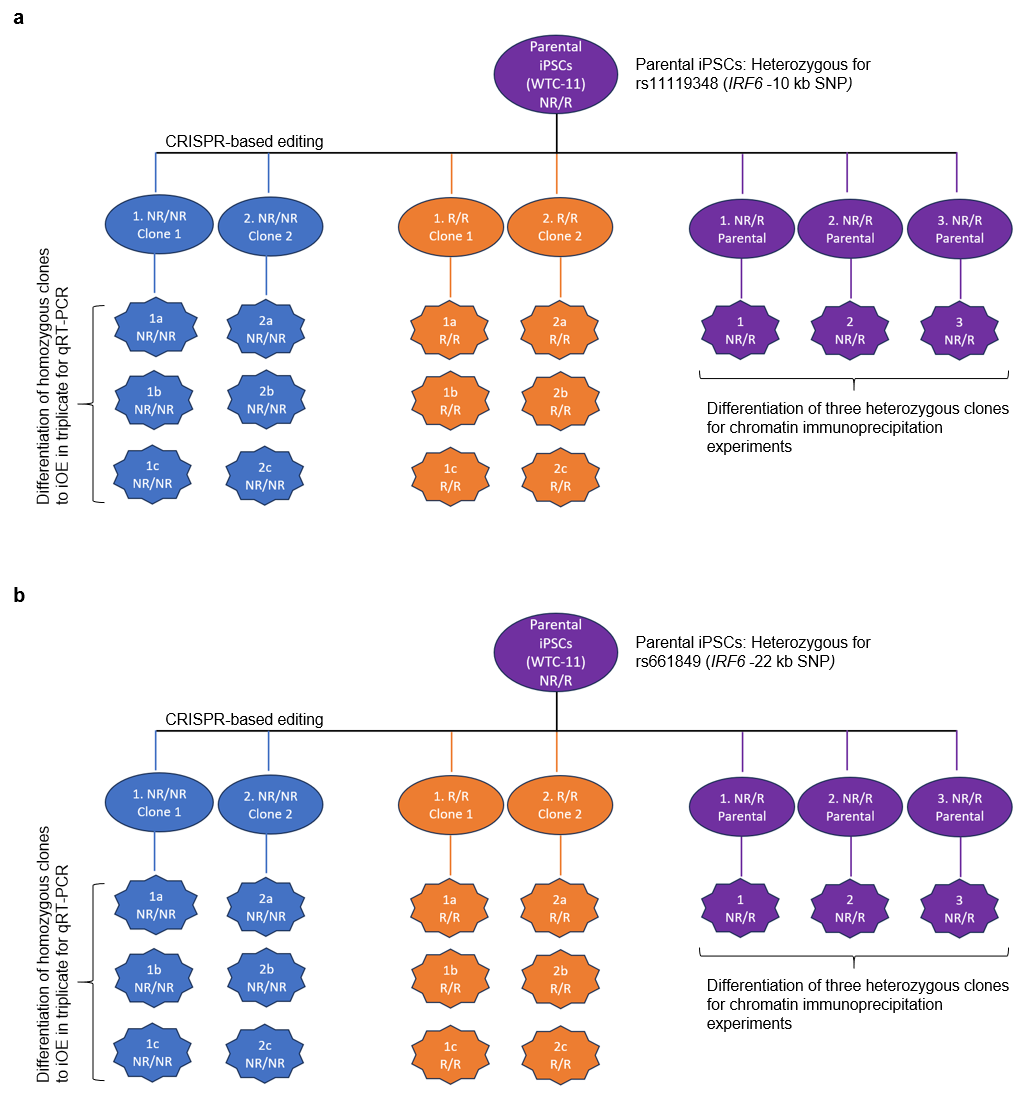

**Supplementary Fig. 13:** Strategy for in vitro cell culture experiments: **a**) Parental induced pluripotent stem cells (iPSCs) WTC-11, heterozygous for rs11119348 (*IRF6* -10 kb SNP) (AC, NR/R), were edited to be homozygous for the non-risk (CC, NR/NR) or the risk allele (AA, R/R) and subjected to a 10-day differentiation protocol to generate induced oral epithelial cells (iOECs). NR, non-risk allele; R, risk allele. **b)** Parental induced pluripotent stem cells (iPSCs) WTC-11, heterozygous for rs661849 (*IRF6* -22 kb SNP) (TC, NR/R), were edited to be homozygous for the non-risk (TT, NR/NR) or the risk allele (CC, R/R) and subjected to a 10-day differentiation protocol to generate induced oral epithelial cells (iOECs). NR, non-risk allele; R, risk allele.

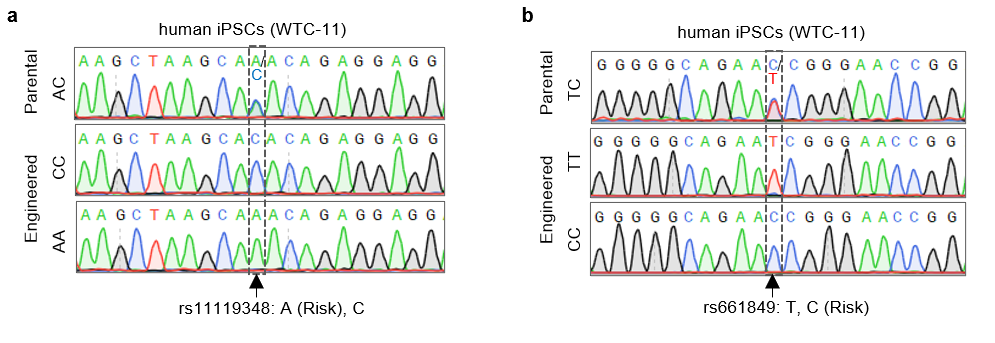

**Supplementary Fig. 14:** Genome of iPSCs engineered to homozygosity for risk and non-risk alleles, each individually, of **a)** the *IRF6* -10 kb SNP (rs11119348), **b)** the *IRF6* -22 kb SNP (rs661849).

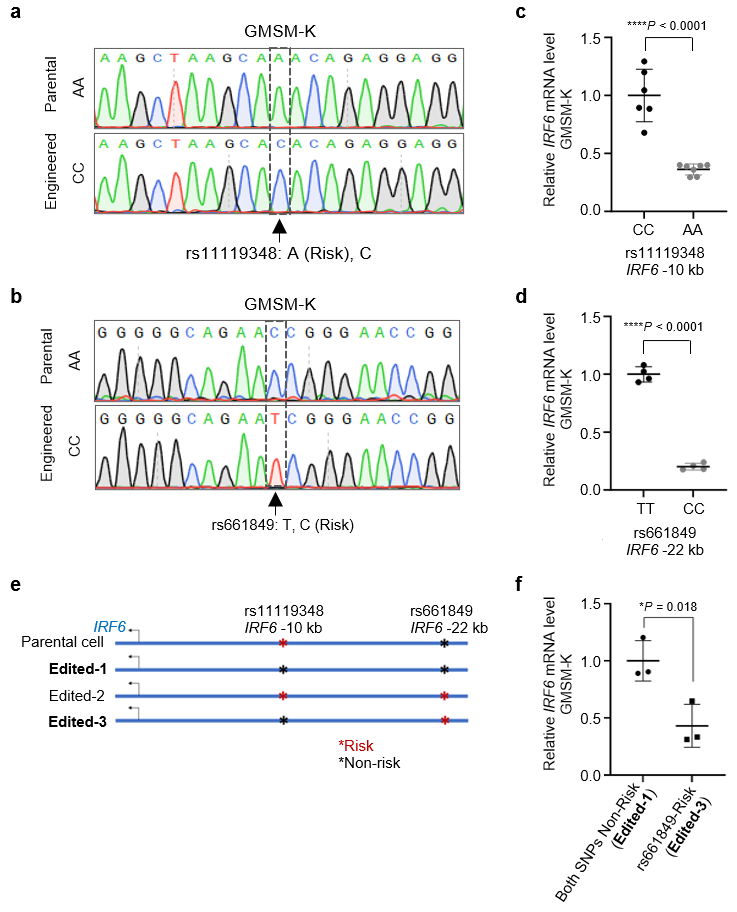

**Supplementary Fig. 15:** *IRF6* RNA expression in engineered GMSM-K: **a** and **b)** GMSM-K engineered to homozygosity for risk and non-risk alleles of the *IRF6* -10 kb SNP and, separately, the *IRF6* -22 kb SNP respectively: **c** and **d)** Scattered dot plot of relative levels of *IRF6* mRNA in GMSM-K harboring **(c)** non-risk (CC) and risk (AA) genotype of rs11119348 and, **(d)** non-risk (TT) and risk (CC) genotype of rs661849 assessed by qRT-PCR. Expression levels of *IRF6* are normalized against *ACTB.* Data are represented as mean ± SD from multiple replicates as indicated in the plot. Statistical significance (*P* value, two-tailed) is determined by Student’s t-test. Please note that GMSM-K harboring non-risk (TT) or risk (CC) genotypes of rs661849 in **b** and **d** harbors the parental risk (AA) genotype of the other SNP (rs11119348) as indicated in panel e. **e)** Engineered genome of GMSM-K harboring non-risk (TT) or risk (CC) genotypes of rs661849 in the background of non-risk (CC) genotype of rs11119348. **f)** Scattered dot plot of relative levels of *IRF6* mRNA in GMSM-K harboring non-risk (TT) and risk (CC) genotype of rs661849 in the background of non-risk (CC) genotype of rs11119348 assessed by qRT-PCR. Expression levels of *IRF6* are normalized against *ACTB.* Data are represented as mean ± SD from multiple replicates as indicated in the plot. Statistical significance (*P* value, two-tailed) is determined by Student’s t-test.

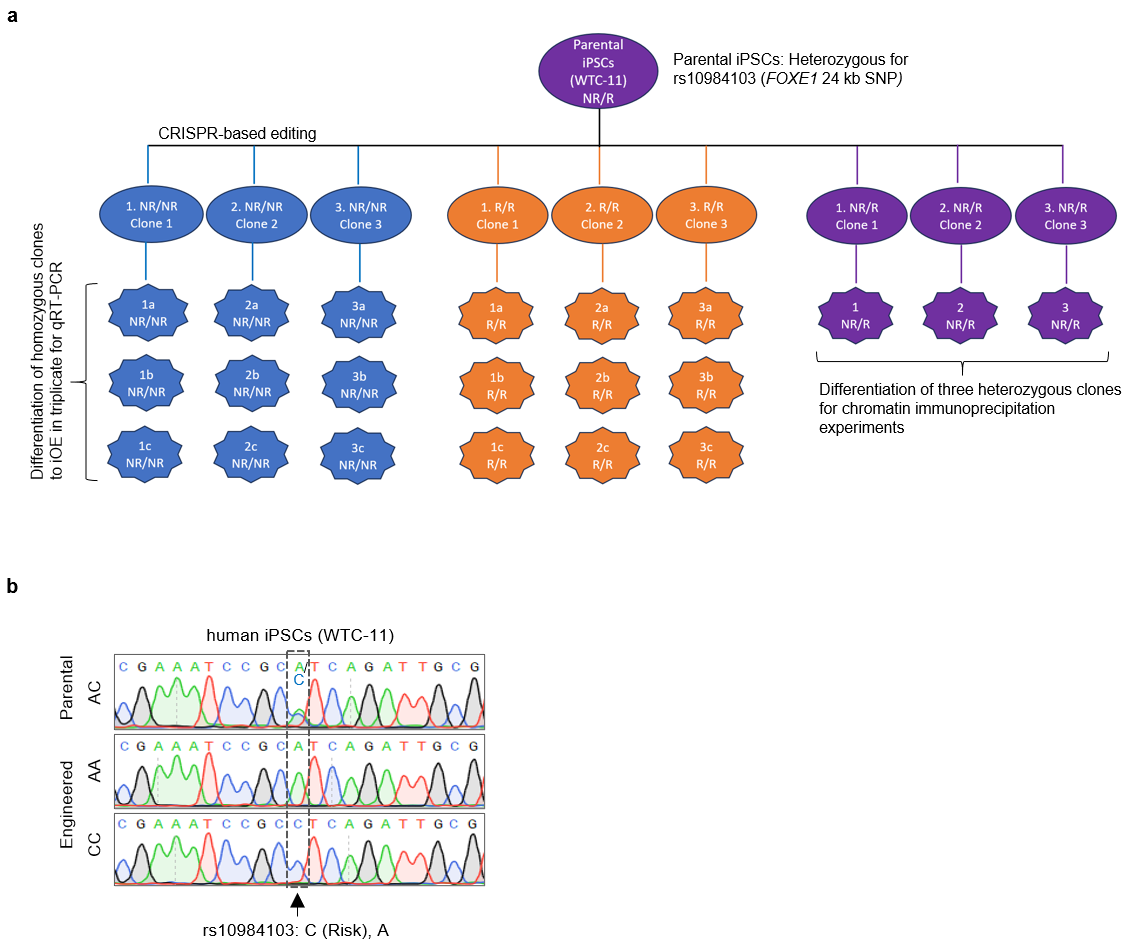

**Supplementary Fig. 16: a)** Strategy for in vitro cell culture experiments: Parental iPSCs WTC-11, heterozygous for rs10984103 (*FOXE1* 24 kb SNP) (AC, NR/R), were edited to be homozygous for the non-risk (AA, NR/NR) or the risk allele (CC, R/R) and subjected to a 10-day differentiation protocol to generate induced oral epithelial cells (iOECs). NR, non-risk allele; R, risk allele. **b)** Genome of iPSCs engineered to homozygosity for risk and non-risk allele, individually for the *FOXE1* 24 kb SNP (rs10984103).

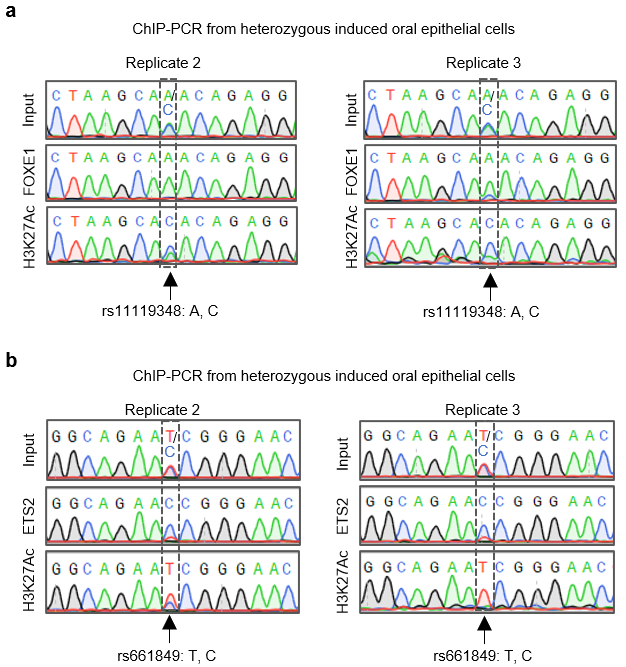

**Supplementary Fig. 17:** Sequencing of **a)** anti-FOXE1 and anti-H3K27Ac ChIP-PCR product of cells heterozygous for rs11119348 from two ChIP replicates and **b)** anti-ETS2 and anti-H3K27Ac ChIP-PCR product of cells heterozygous for rs661849 from two ChIP replicates.

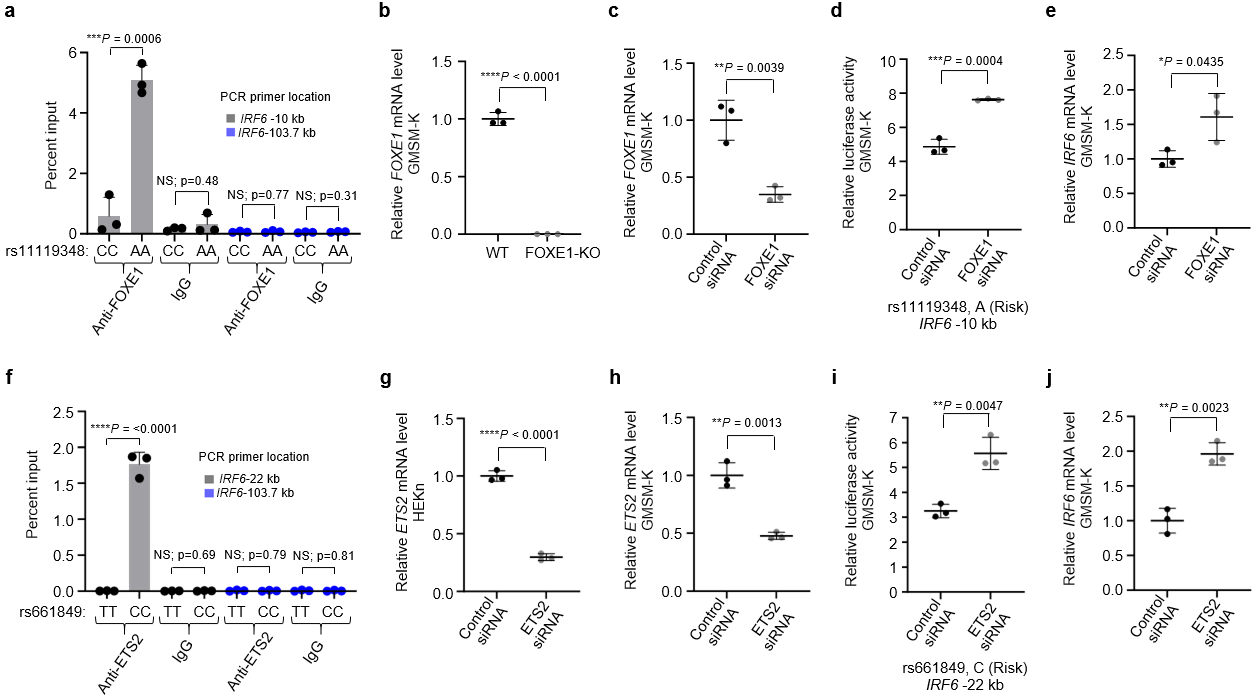

**Supplementary Fig. 18: a)** Percent input identified by ChIP-qPCR for anti-FOXE1 in GMSM-K harboring non-risk (CC) and risk (AA) genotype of rs11119348 using primers specific to the *IRF6* -10 kb enhancer site or, as a negative control, to a region 103.7 kb upstream of the *IRF6* transcription start site that lacked ATAC-Seq and H3K27Ac ChIP-Seq signals in HIOEC or NHEK. Error bars refer to three ChIP replicates and expressed as mean ± SD. Statistical significance (*P* value, two-tailed) is determined by Student’s *t*-test. NS, non-significant. **b** and **c)** Scattered dot plot of relative levels of *FOXE1* mRNA **(b)** in WT and *FOXE1* KO GMSM-K **(c)** or control siRNA and FOXE1 targeting siRNA transfected GMSM-K assessed by qRT-PCR. Expression levels of *FOXE1* are normalized against *ACTB.* Data are represented as mean ± SD from three replicates. Statistical significance (*P* value, two-tailed) is determined by Student’s *t*-test. For the experiments (Supplementary Figs. 18b-e) the GMSM-K cells are homozygous risk (AA) for the *IRF6* -10 kb SNP. **d)** Scattered dot plot of relative luciferase activity for longer construct with risk allele (A) of rs11119348 in GMSM-K transfected with control siRNA or FOXE1 targeting siRNA. Data are represented as mean ± SD from three independent experiments. Statistical significance (*P* value, two-tailed) is determined by Student’s *t*-test. **e)** Scattered dot plot of relative levels of *IRF6* mRNA in control siRNA and FOXE1 targeting siRNA transfected GMSM-K assessed by qRT-PCR. Expression levels of *IRF6* are normalized against *ACTB.* Data are represented as mean ± SD from three replicates. Statistical significance (*P* value, two-tailed) is determined by Student’s *t*-test. **f)** Percent input identified by ChIP-qPCR for anti-ETS2 in GMSM-K harboring non-risk (TT) and risk (CC) genotype of rs661849 using primers specific to the *IRF6* -22 kb enhancer site or, as a negative control, to a region 103.7 kb upstream of the *IRF6* transcription start site that lacked ATAC-Seq and H3K27Ac ChIP-Seq signals in HIOEC or NHEK. Error bars refer to 3 ChIP replicates and expressed as mean ± SD. Statistical significance (*P* value, two-tailed) is determined by Student’s *t*-test. NS, non-significant. **g, h)** Scattered dot plot of relative levels of *ETS2* mRNA in control siRNA and ETS2 targeting siRNA transfected (**g)** (HEKn) or **(h)** GMSM-K assessed by qRT-PCR. Expression levels of *ETS2* are normalized against *ACTB.* Data are represented as mean ± SD from three replicates. Statistical significance (*P* value, two-tailed) is determined by Student’s *t*-test. SiRNA transfection experiments are performed in HEKn heterozygous (TC) for the *IRF6* -22 kb SNP (Supplementary Figs. 18g) or in GMSM-K harboring homozygous risk genotype (CC) for the *IRF6* -22 kb SNP (Supplementary Figs. 18h-j). **i)** Scattered dot plot of relative luciferase activity for longer construct with risk allele (C) of rs661849 in GMSM-K transfected with control siRNA or ETS2 targeting siRNA. Data are represented as mean ± SD from three independent experiments. Statistical significance (*P* value, two-tailed) is determined by Student’s *t*-test. **j)** Scattered dot plot of relative levels of *IRF6* mRNA in control siRNA and ETS2 targeting siRNA transfected GMSM-K assessed by qRT-PCR. Expression levels are normalized against *ACTB.* Data are represented as mean ± SD from three replicates. Statistical significance (*P* value, two-tailed) is determined by Student’s *t*-test.

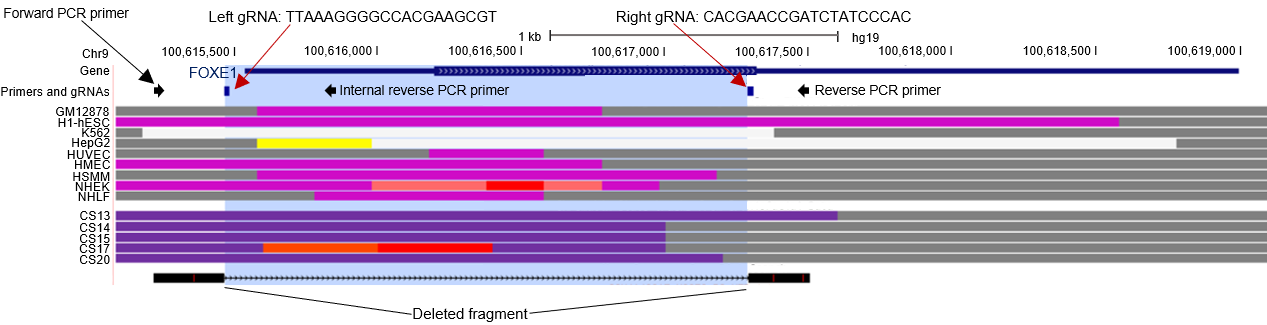

**Supplementary Fig. 19:** *FOXE1* KO GMSM-K: Browser view of the human genome, GRCh37/hg19, focused on *FOXE1* gene. **Primers and gRNAs**, a pair of guide RNAs (marked with a block) and primers (marked with arrows) used to generate *FOXE1* KO cells. Primer sequences are provided in Supplementary Table 20b. Next two track and color codes are similar as in Fig. 2. The blue region highlights the deleted fragment after CRISPR/Cas9 targeting.

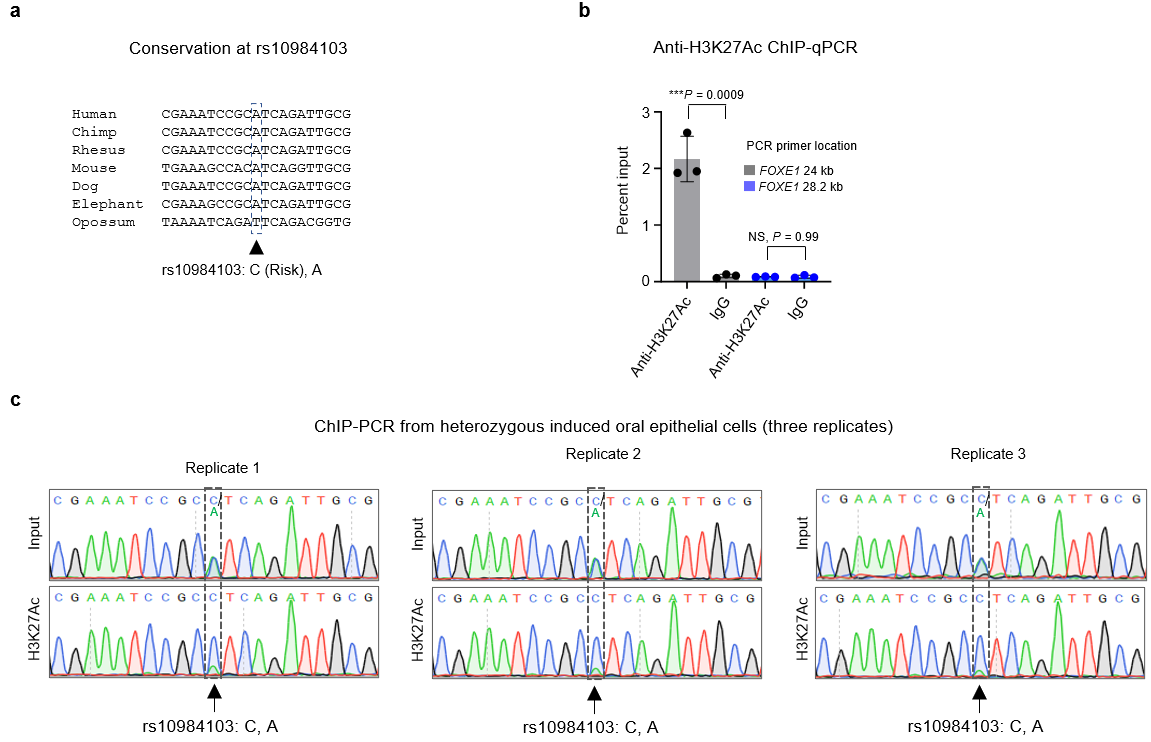

**Supplementary Fig. 20: Risk allele of *FOXE1* 24 kb SNP (****rs10984103) increases enhancer activity. a)** Alignment of the rs10984103 variant site in different species (https://genome.ucsc.edu). At this SNP, the risk allele ‘C’ has a higher frequency than the non-risk and referent allele ‘A’ in all populations. **b)** Percent input identified by ChIP-qPCR for anti-H3K27Ac in iOE cells heterozygous for rs10984103 using primers specific to the *FOXE1* 24 kb enhancer site or, as a negative control, to a region 28.2 kb downstream of the *FOXE1* transcription start site that lacked ATAC-Seq and H3K27Ac ChIP-Seq signals in HIOEC or NHEK. Error bars refer to three ChIP replicates and expressed as mean ± SD. Statistical significance (*P* value, two-tailed) is determined by Student’s *t*-test. NS, non-significant. **c)** Sequencing of anti-H3K27Ac ChIP-PCR product of iOE cells heterozygous for rs10984103 from three replicates.

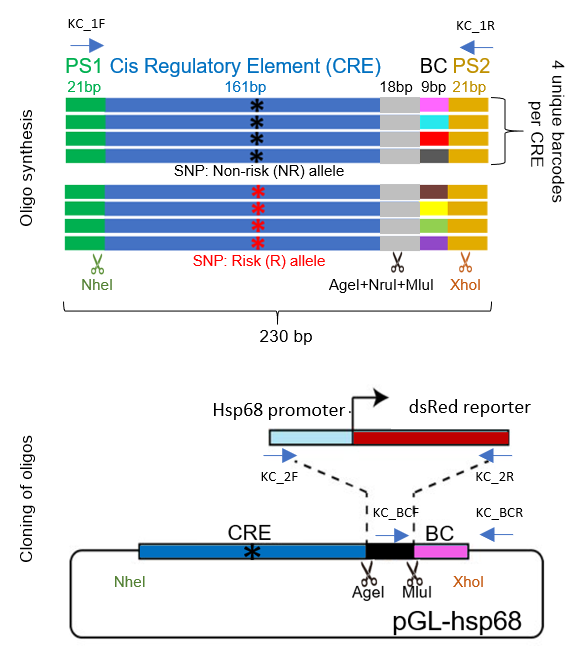

**Supplementary Fig. 21:** Details of MPRA library construction. PS represents priming sequences; BC represents barcode. KC represents different set of primers used in the MPRA library process (see method section and Supplementary Table 18 for details).

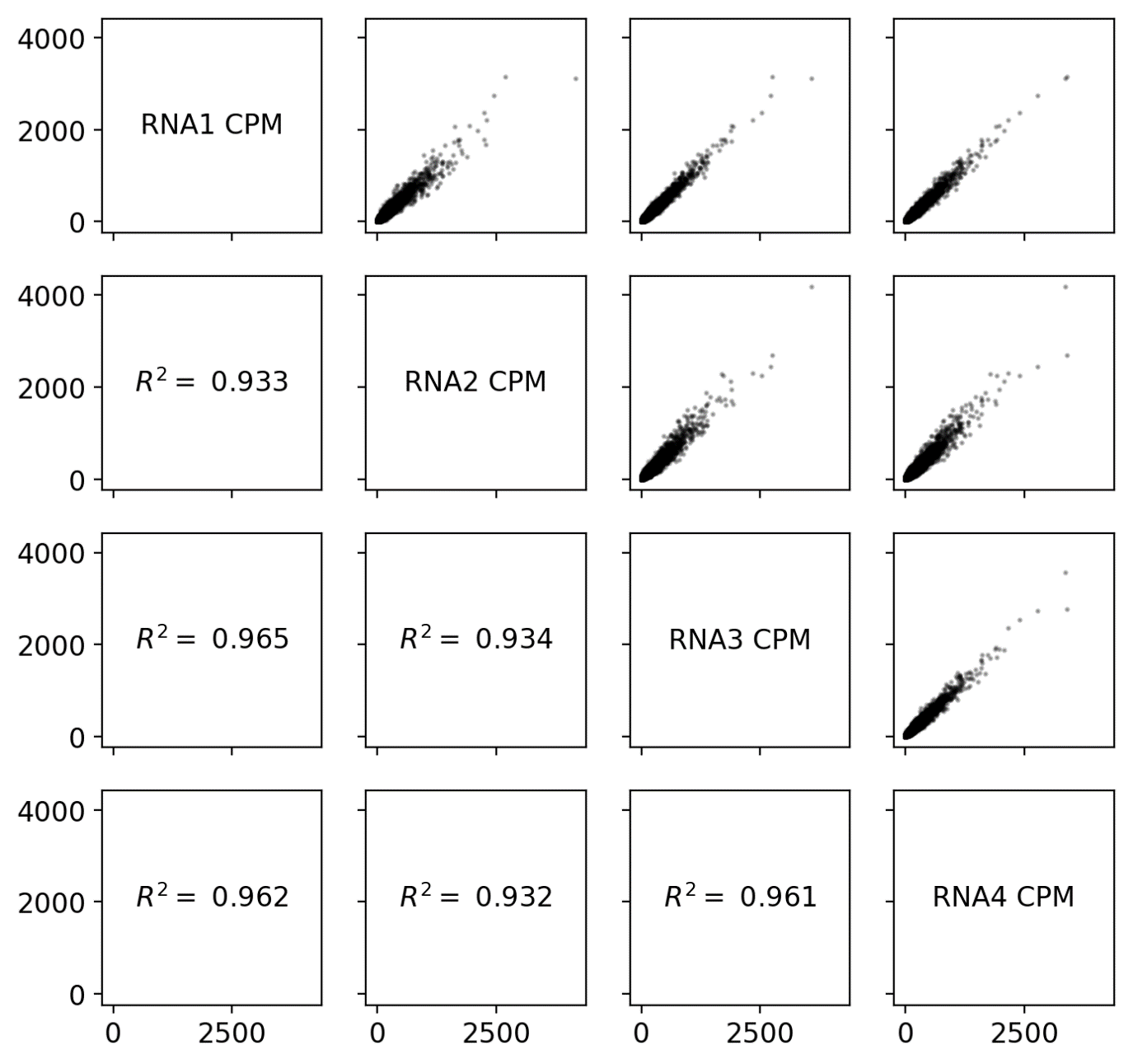

**Supplementary Fig. 22:** Correlation plot showing distribution of RNA barcode counts across four RNA replicates.

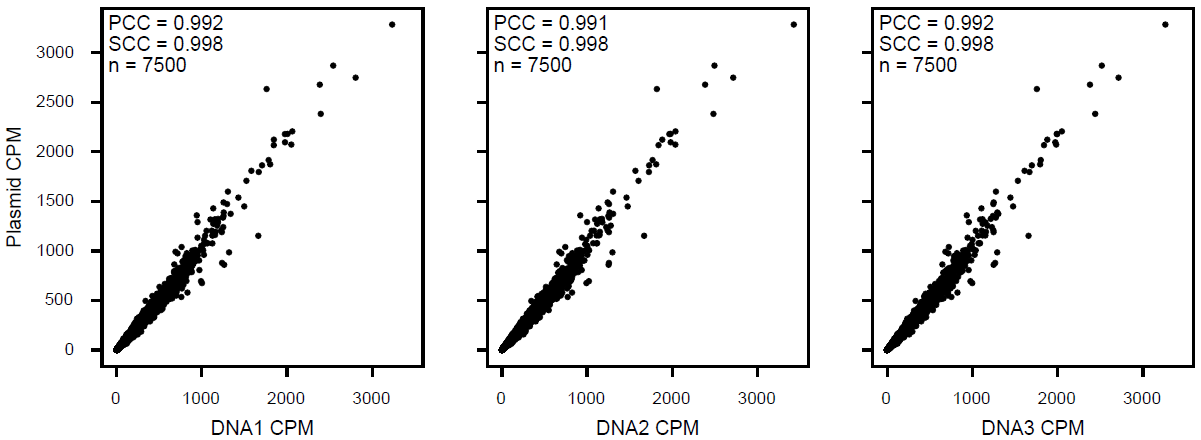

**Supplementary Fig. 23:** Correlation plot showing DNA barcode counts (post-transfection) vs plasmid DNA barcode counts (pre-transfection).

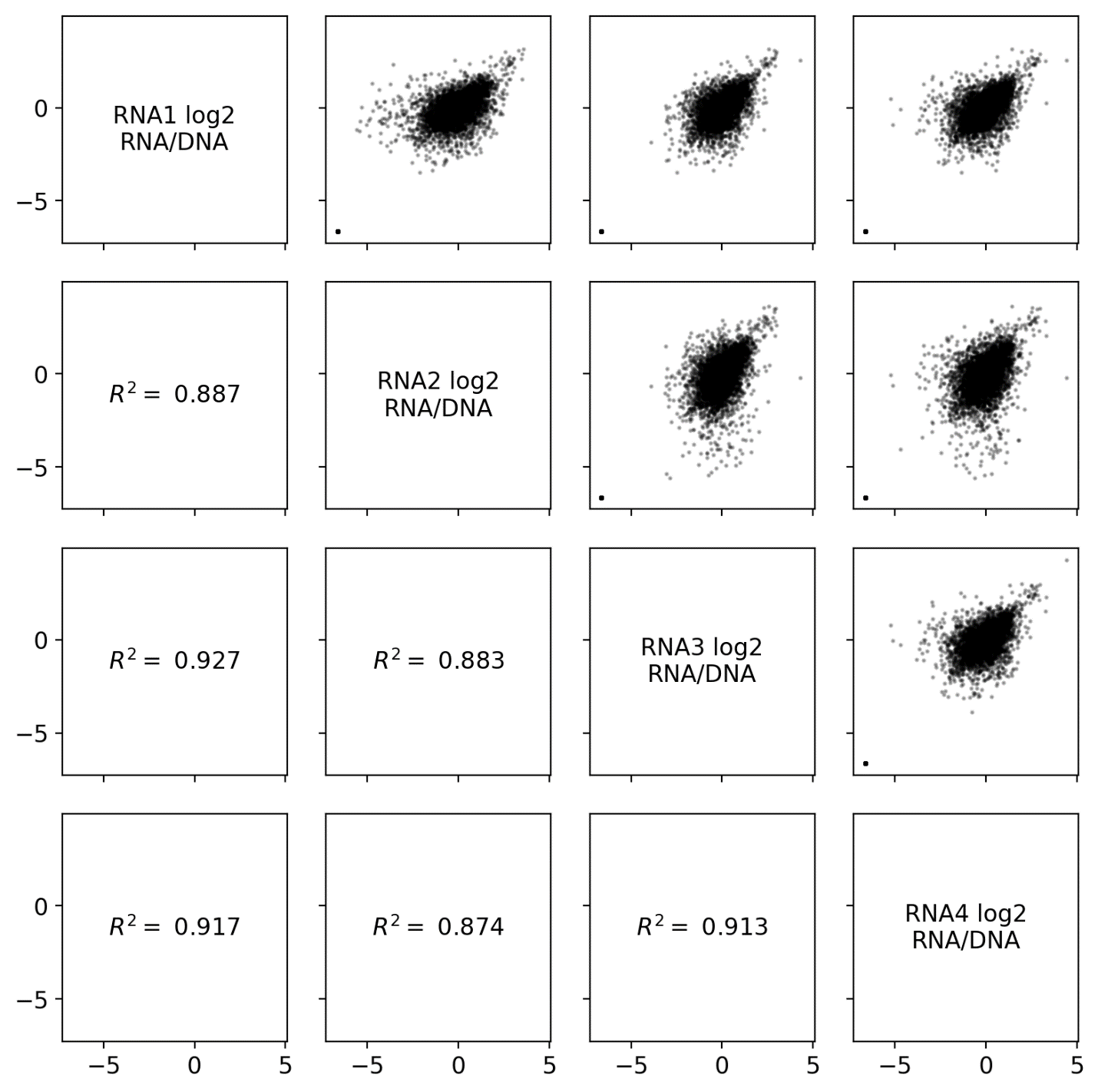

**Supplementary Fig. 24:** Correlation plot showing RNA barcode counts normalized to plasmid DNA barcode counts (pre-transfection) across four replicates.
